# A transgenic zebrafish line for in vivo visualisation of neutrophil myeloperoxidase

**DOI:** 10.1101/456541

**Authors:** Kyle D. Buchan, Tomasz K. Prajsnar, Nikolay V. Ogryzko, Nienke W.M. de Jong, Michiel van Gent, Julia Kolata, Simon J. Foster, Jos A.G. van Strijp, Stephen A. Renshaw

**Affiliations:** The Bateson Centre and Department of Infection, Immunity and Cardiovascular Disease, University of Sheffield, Western Bank, Sheffield, S10 2TN, UK.; Department of Medical Microbiology, University Medical Center Utrecht, Utrecht University, Utrecht, The Netherlands.; Centre for Inflammation Research, University of Edinburgh, Edinburgh, EH16 4TJ, United Kingdom.; Department of Molecular Biology and Biotechnology, University of Sheffield, Western Bank, Sheffield, S10 2TN, UK.

## Abstract

The neutrophil enzyme myeloperoxidase (MPO) is a major enzyme made by neutrophils to generate antimicrobial and immunomodulatory compounds, notably hypochlorous acid (HOCl), amplifying their capacity for destroying pathogens and regulating inflammation. Despite its roles in innate immunity, the importance of MPO in preventing infection is unclear, as individuals with MPO deficiency are asymptomatic with the exception of an increased risk of candidiasis. Dysregulation of MPO activity is also linked with inflammatory conditions such as atherosclerosis, emphasising a need to understand the roles of the enzyme in greater detail. Consequently, new tools for investigating granular dynamics *in vivo* can provide useful insights into how MPO localises within neutrophils, aiding understanding of its role in preventing and exacerbating disease. The zebrafish is a powerful model for investigating the immune system *in vivo*, as it is genetically tractable, and optically transparent.

To visualise MPO activity within zebrafish neutrophils, we created a genetic construct that expresses human MPO as a fusion protein with a C-terminal fluorescent tag, driven by the neutrophil-specific promoter *lyz*. After introducing the construct into the zebrafish genome by Tol2 transgenesis, we established the *Tg(lyz:Hsa.MPO-mEmerald,cmlc2:EGFP)sh496* line, and confirmed transgene expression in zebrafish neutrophils. We observed localisation of MPO-mEmerald within a subcellular location resembling neutrophil granules, mirroring MPO in human neutrophils. In Spotless (*mpx^NL144^*) larvae - which express a non-functional zebrafish myeloperoxidase - the MPO-mEmerald transgene does not disrupt neutrophil migration to sites of infection or inflammation, suggesting that it is a suitable line for the study of neutrophil granule function.

We present a new transgenic line that can be used to investigate neutrophil granule dynamics *in vivo* without disrupting neutrophil behaviour, with potential applications in studying processing and maturation of MPO during development.

## Introduction

The enzyme Myeloperoxidase (MPO) enhances the microbicidal potential of neutrophils by converting hydrogen peroxide (H_2_O_2_) into the highly toxic antimicrobial compound hypochlorous acid (HOCl) (1), and by forming radicals by oxidating substrates including phenols, nitrate and tyrosine residues (2). MPO is located in the primary granules of neutrophils, which deliver MPO and other bactericidal compounds to invading pathogens by fusing with phagocytic vesicles, accelerating pathogen destruction. MPO is the most abundant protein in the primary granules of human neutrophils (3), and consequently neutrophils are able to produce high levels of HOCl to deliver a potent antimicrobial response that is capable of killing a broad variety of major pathogens (4–6).

Importantly, due to the activity of upstream NADPH oxidase, the phagocytic vacuole is thought to be relatively alkaline (~pH 9), and under such conditions MPO activity may be less efficient (7) than other neutrophil enzymes (8). MPO activity appears to be context-dependent, particularly during phagocytosis of large structures such as fungal hyphae (9) or bacterial biofilms (10). In these cases, the phagocytic vacuole does not fully close (11), causing MPO to act at the acidic pH of sites of inflammation (~pH 6) (12), at which it can function normally. This observation is supported by the fact that MPO is thought to play a role in the generation of neutrophil extracellular traps (NETs) (13), which are often induced in response to large targets (14). While the precise site of action of MPO is uncertain, it is clear that it plays a role in antimicrobial defence, as pathogens produce specific virulence factors targeting it (15).

Beyond its role in bolstering the antimicrobial defence, MPO is also an important regulator of inflammation. The arrival of neutrophils at the wound site marks the initial steps of the anti-inflammatory response, as MPO is delivered to the wound site to consume H_2_O_2_ and reduce inflammatory signalling (16,17). There is also a link between aberrant MPO activity and inflammatory conditions: overactivity is associated with cardiovascular disease, multiple sclerosis and glomerulonephritis (18–20), while MPO deficiency has been implicated in pulmonary fibrosis and atherosclerosis (21,22), highlighting its critical role in immune homeostasis. MPO deficiency is a relatively common condition affecting 1 in every 2,000-4,000 people across Europe and North America (23), with no major health risks apart from a susceptibility to *Candida albicans* infections (24). This observation is in stark contrast to people with chronic granulomatous disease (CGD), who lack a working NADPH oxidase. Those with CGD are unable to generate an effective phagocytic environment capable of destroying microbes (25,26). Unlike MPO deficiency, those with CGD experience frequent life-threatening infections from a wide range of pathogens (27), and consequently, the role of MPO is less clear when observed in the context of other oxidative enzymes and compounds. Further studies are required to understand the complex roles of MPO in the immune system.

The zebrafish is a powerful model for studying physiology and pathology *in vivo* and has been used to model many important conditions ranging from neurodegenerative disorders such as Alzheimer’s disease (28), to cancers including melanoma (29) and leukaemia (30). They are optically transparent, making them amenable to imaging studies and produce high numbers of offspring, which permits the application of high-throughput approaches. Another major advantage of the zebrafish is their genetic tractability, facilitating the introduction of large genetic constructs into the genome, often expressing fluorescent proteins driven by tissue-specific promoters (31). Several studies have utilised these features to create transgenic lines labelling macrophages (32) and neutrophils (33) to image the innate immune response during infection (34) and inflammation (35).

MPO can be measured using a variety of cytochemical and cytometry-based approaches (36), however there are relatively few tools that allow granular MPO to be visualised *in vivo* and in real time. Mouse models that permit imaging of neutrophil granules and MPO do exist (20,37), however murine MPO lacks several transcription factor binding domains (38), and is expressed at 1/10 the level found in human neutrophils (39), raising concerns over whether a murine model can fully represent human MPO.

In this study, we have generated a transgenic zebrafish line expressing fluorescently-labelled human MPO in zebrafish neutrophils, as a tool towards investigating the roles of MPO during infection and inflammation. The transgene for the MPO-mEmerald fusion protein (*lyz:*Hsa.MPO-mEmerald) was successfully expressed in zebrafish neutrophils and the resulting protein appears to be trafficked to granules, recapitulating expression of MPO in human neutrophils. Additionally, we showed that the MPO-mEmerald enzyme does not disrupt neutrophil recruitment to sites of injury and infection. In the future, *Tg(lyz:Hsa.MPO-mEmerald,cmlc2:EGFP)sh496* zebrafish may prove to be a useful tool for investigating MPO and imaging granular dynamics *in vivo* and in real-time.

## Methods

### Zebrafish Husbandry

Zebrafish (*Danio rerio*) were raised and maintained under the Animals (Scientific Procedures) Act 1986 using standard protocols (40). Adult zebrafish were hosted in UK Home Office-approved aquaria at the Bateson Centre, University of Sheffield, and kept under a 14/10 light/dark regime at 28°C.

### Cloning the *Tg(lyz:Hsa.MPO-mEmerald,cml2:EGFP)sh496* line

The plasmid used for introducing the transgene into the zebrafish genome (pDestTol2CG2 *lyz:*MPO-mEmerald *cmlc2*:EGFP) was created by Gateway cloning (31). Briefly, a gateway vector p5E-MCS containing 6.6kb of the lysozyme C promoter (33) was used to drive neutrophil-specific expression,. The MPO-mEmerald gene was incorporated into an expression vector by first digesting the mEmerald-MPO-N-18 plasmid (Addgene plasmid #54187, Dr. Michael Davidson’s lab), and ligating the MPO-mEmerald fusion protein gene into the multiple cloning site vector pME MCS to generate the middle-entry vector, pME MCS MPO-mEmerald. The final construct was created by an LR reaction combining a 5’ vector containing the *lyz* promoter, the middle entry vector pME MPO-mEmerald, a 3’ vector containing a polyadenylation site, and the destination vector pDestTol2CG2.

### Generation of the *Tg(lyz:nfsB-mCherry)sh260* line

As for the MPO-mEmerald construct, the *lyz* promoter in the gateway vector p5E-MCS was used to drive neutrophil-specific expression. For red fluorescence, mCherry was produced as a fusion protein with the nitroreductase gene *nfsB* (primers F: 5’ GGG GAC AAG TTT GTA CAA AAA AGC AGG CTG CAT GGA TAT CAT TTC TGT CGC CTT, R: 5’ GGG GAC CAC TTT GTA CAA GAA AGC TGG GTC GGT CCA CTT CGG TTA AGG TGA TGT T), which also permits conditional ablation of cells upon addition of metronidazole (41), and cloned into middle entry (pME-nfsB) and 3’ entry vectors (p3E-mCherry). The final lyz:nfsB-mCherry construct was created by recombination of p5E-lyz, pME-nfsB, and p3E-mCherry, and pDestTol2pA2.

### Microinjection of *lyz:*MPO-mEmerald Construct DNA

Construct DNA of the donor plasmid pDestTol2CG2 *lyz:*MPO-mEmerald *cmlc2*:EGFP or pDestTol2pA2 *lyz:*nfsB-mCherry was injected into zebrafish embryos at the one-cell stage with 10ng/µl of Tol2 transposase RNA, according to published protocols (40).

### TSA staining and colocalisation experiments

3dpf *Tg(lyz:Hsa.MPO-mEmerald,cmlc2:EGFP)sh496* larvae were fixed in ice-cold 4% (w/v) paraformaldehyde (PFA) in PBS-TX (PBS supplemented with 0.5 % of Triton X-100) overnight at 4°C. Fixed embryos were washed in PBS-TX twice. Peroxidase activity was detected by incubation in 1:50 Cy5-TSA:amplification reagent (PerkinElmer, Waltham, MA) in the dark for 10 min at 28°C followed by extensive washing in PBS-TX.

### Zebrafish Tailfin Transection

*Tg(lyz:Hsa.MPO-mEmerald,cmlc2:EGFP)sh496* zebrafish at 3 days post-fertilisation were anaesthetised by immersion in E3 supplemented with 4.2% Tricaine and complete transection of the tail was performed with a sterile scalpel. For imaging of larvae, a Nikon® custom-build wide-field microscope was used: Nikon® Ti-E with a CFI Plan Apochromat λ 10X, N.A.0.45 objective lens, a custom built 500 μm Piezo Z-stage (Mad City Labs, Madison, WI, USA) and using Intensilight fluorescent illumination with ET/sputtered series fluorescent filters 49002 and 49008 (Chroma, Bellow Falls, VT, USA) was used. Analysis was performed using Nikon’s® NIS Elements software package.

### Bacterial Culture Preparation

To prepare a liquid overnight culture of *S. aureus*, 5ml of BHI broth medium (Oxoid) was inoculated with a colony of *S. aureus* strain USA300, and incubated at 37°C overnight with shaking. To prepare *S. aureus* for injection, 50ml of BHI media was inoculated with 500µl of overnight culture and incubated for roughly 2 hours at 37°C with shaking. The OD_600_ of each culture was measured and 40ml of the remaining culture harvested by centrifugation at 4,500g for 15 minutes at 4°C. The pellet was then resuspended in a volume of PBS appropriate to the bacterial dose required. Once the pellets were resuspended they were then kept on ice until required.

### Spotless (*mpx^NL144^*) Fish and Sudan Black Staining

The Spotless (*mpx^NL144^*) mutant line contains a C to T mutation at nucleotide 1126 of the *mpx* RefSeq mRNA sequence (NM_212779), resulting in a premature stop codon (42). A detailed protocol of Sudan Black B staining can be found in Supplementary File 1.

### Microscopy of Neutrophil Granules

Microscopy of neutrophil granules in *Tg(lyz:Hsa.MPO-mEmerald,cmlc2:EGFP)sh496* larvae was performed using a Zeiss® Axiovert LSM 880 Airyscan confocal microscope with 63x Plan Apochromat oil objective (NA 1.4). Cells were illuminated with a 488 nm argon laser and/or a 561 nm diode laser. Images were processed using the Zeiss® microscope software and analysed using Zen Black.

### Statistics

All data were analysed (Prism 7.0, GraphPad Software, San Diego, CA, USA) using a two-way ANOVA with Bonferroni post-test to adjust for multiple comparisons.

## Results

### Creation of a transgenic zebrafish expressing human myeloperoxidase

To create a transgenic zebrafish that expresses a fluorescently-tagged human myeloperoxidase (MPO), we created a genetic construct using Gateway cloning that contains the MPO gene with a C-terminal fusion of the fluorescent protein mEmerald, driven by the neutrophil-specific promoter *lyz* (Figure 1A). After the construct was successfully assembled, it was introduced into the zebrafish genome by Tol2-mediated transgenesis. To verify expression in neutrophils, a second transgenic line was created using a construct expressing mCherry under the *lyz* promoter, *Tg(lyz:nfsB-mCherry)sh260*. Successful expression of MPO-mEmerald in zebrafish neutrophils was confirmed by inducing transgenesis in *Tg(lyz:nfsB-mCherry)sh260* fish (Figure 1CD). Injected larvae were then screened at 3 days post fertilisation (dpf) for mEmerald expression and colocalisation with mCherry expression. Figure 1CD shows double-transgenic neutrophils expressing both mEmerald and mCherry in the primary haematopoietic tissue of the zebrafish larvae, the caudal haematopoietic tissue (CHT) (indicated in Figure 1B) (43). This observation confirms that the construct is successfully expressed and suggests that it co-localises with zebrafish neutrophils. We also noted that in double-transgenic neutrophils, there appeared to be a differential subcellular localisation between mEmerald and mCherry signal, with mCherry localised to areas with no visible mEmerald signal (Figure 1D).

**Figure 1.**
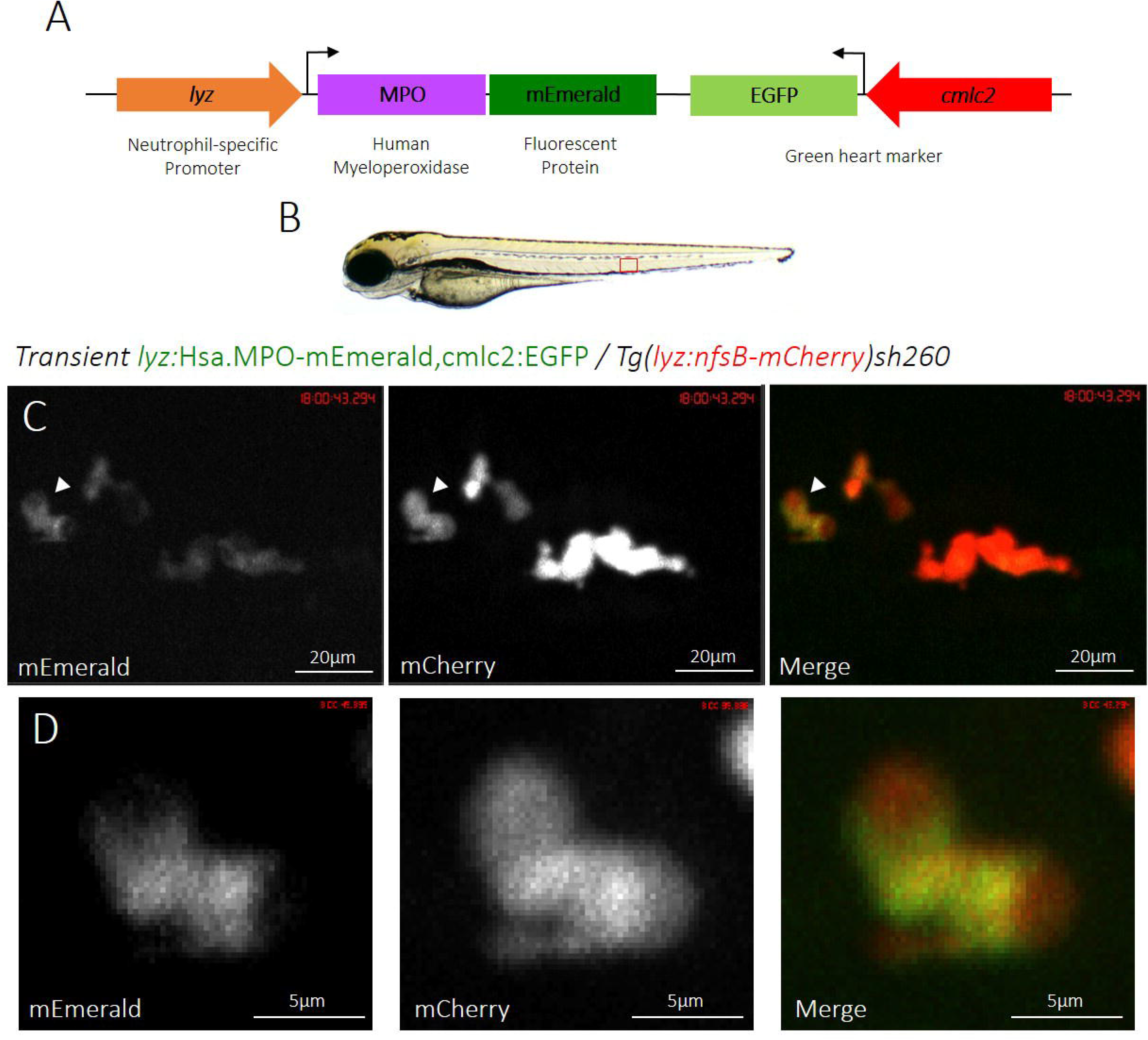
Transient expression of the *lyz:*MPO-mEmerald transgene labels zebrafish neutrophils **A)** Schematic of the *lyz*:MPO-mEmerald *cmlc2*:EGFP construct, which includes the neutrophil-specific promoter (*lyz*), the MPO gene with a C-terminal fluorescent tag (MPO-mEmerald) and a green heart marker to aid optimisation of transgenesis (*cmlc2:*EGFP). **B)** A zebrafish larva at 3 days post fertilisation (dpf), with the caudal haematopoietic tissue (CHT) indicated by the red box. **C)** The CHT of a double-transgenic *Transient lyz:*MPO-mEmerald,*cmlc2*:EGFP; *Tg(lyz:nfsB-mCherry)sh260* larva with a population of neutrophils expressing both mEmerald and mCherry. The white arrowhead indicates the neutrophil enlarged below. **D)** Enlarged view of a neutrophil expressing mEmerald and mCherry.

### MPO-mEmerald is stably expressed in zebrafish neutrophils

To secure adult zebrafish with stable germline integrations of the *lyz:*MPO-mEmerald transgene, larvae that transiently expressed the transgene were identified, raised and outcrossed to determine whether the transgene was inherited by their offspring. An adult that produced larvae with a cell population labelled with mEmerald was identified and its progeny raised to produce fish stably expressing the MPO transgene, with the designation *Tg(lyz:Hsa.MPO-mEmerald,cmlc2:EGFP)sh496*. To verify whether the *lyz:*MPO-mEmerald transgene was expressed in neutrophils of stably transgenic fish, they were crossed to the red neutrophil reporter line *Tg(lyz:nfsB-mCherry)sh260*, and screened for any co-expression of fluorescent proteins. Both transgenes were expressed in neutrophils throughout the CHT (Figure 2), demonstrating that *lyz:*MPO-mEmerald is expressed in zebrafish neutrophils in stably transgenic larvae.

**Figure 2.**
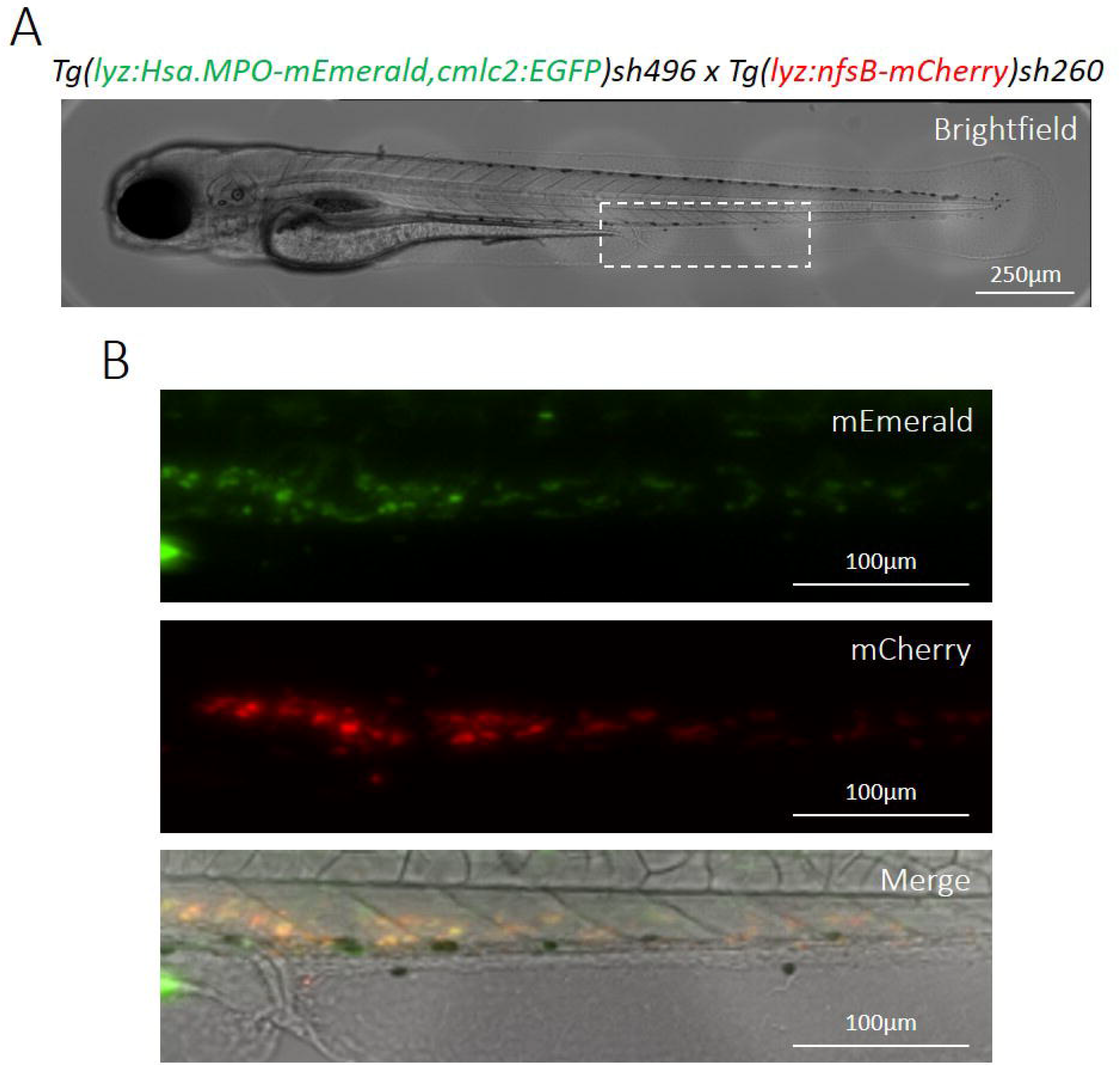
Transgenic *Tg(lyz:Hsa.MPO-mEmerald,cmlc2:EGFP)sh496* zebrafish stably express the transgene in zebrafish neutrophils **A)** A brightfield view of a double-transgenic *Tg(lyz:Hsa.MPO- mEmerald,cmlc2:EGFP)sh496*; *Tg(lyz:nfsB-mCherry)sh260* zebrafish larva at 3dpf. The dashed white box indicates the enlarged region shown in B). **B)** mEmerald and mCherry expression in the CHT of the larva shown in A).

To further confirm that MPO-mEmerald positive cells are neutrophils, we performed TSA-staining on fixed larvae at 3dpf (Figure 4), to stain specific endogenous peroxidase activity in zebrafish neutrophils (44). An average of 80% of the cells observed in the CHT are positive for both MPO-mEmerald and TSA, suggesting that the transgene specifically labels neutrophils in the larva – the staining of neutrophils with TSA is incomplete accounting for the less than 100% co-staining.

### MPO-mEmerald is trafficked to a subcellular location

As MPO is located in the primary granules of neutrophils prior to delivery to the phagosome (1), we wished to determine whether the *lyz:*MPO-mEmerald transgene recapitulates MPO expression in human neutrophils. To investigate the intracellular localisation of the MPO transgene, *Tg(lyz:Hsa.MPO-mEmerald,cmlc2:EGFP)sh496* fish were outcrossed to *Tg(lyz:nfsB-mCherry)sh260* fish, and at 3dpf the double-transgenic larvae were imaged in high detail using an Airyscan confocal microscope. Both transgenes are expressed in the same cells, with MPO-mEmerald localising with a granular subcellular distribution (Figure 3), suggesting that the MPO-mEmerald fusion protein is trafficked to and packaged within neutrophil granules. High-speed imaging of a double-transgenic *Tg(lyz:Hsa.MPO-mEmerald,cmlc2:EGFP)sh496; Tg(lyz:nfsB-mCherry)sh260* larva shows that the intracellular MPO-mEmerald signal is highly dynamic (Supplemental movie 1), and resembles the Brownian motion that would be observed in primary neutrophil granules. Additionally, we also analysed colocalisation of the MPO-mEmerald signal with the location of TSA histochemical staining in neutrophils (Figure 5). There was 70% colocalisation between MPO-mEmerald and endogenous peroxidase activity (the MPO transgene does not have peroxidase activity – see below). This suggests that MPO-mEmerald is expressed in neutrophil granules, with the differences in co-localisation due to inactive, unprocessed forms of MPO-mEmerald.

**Figure 3.**
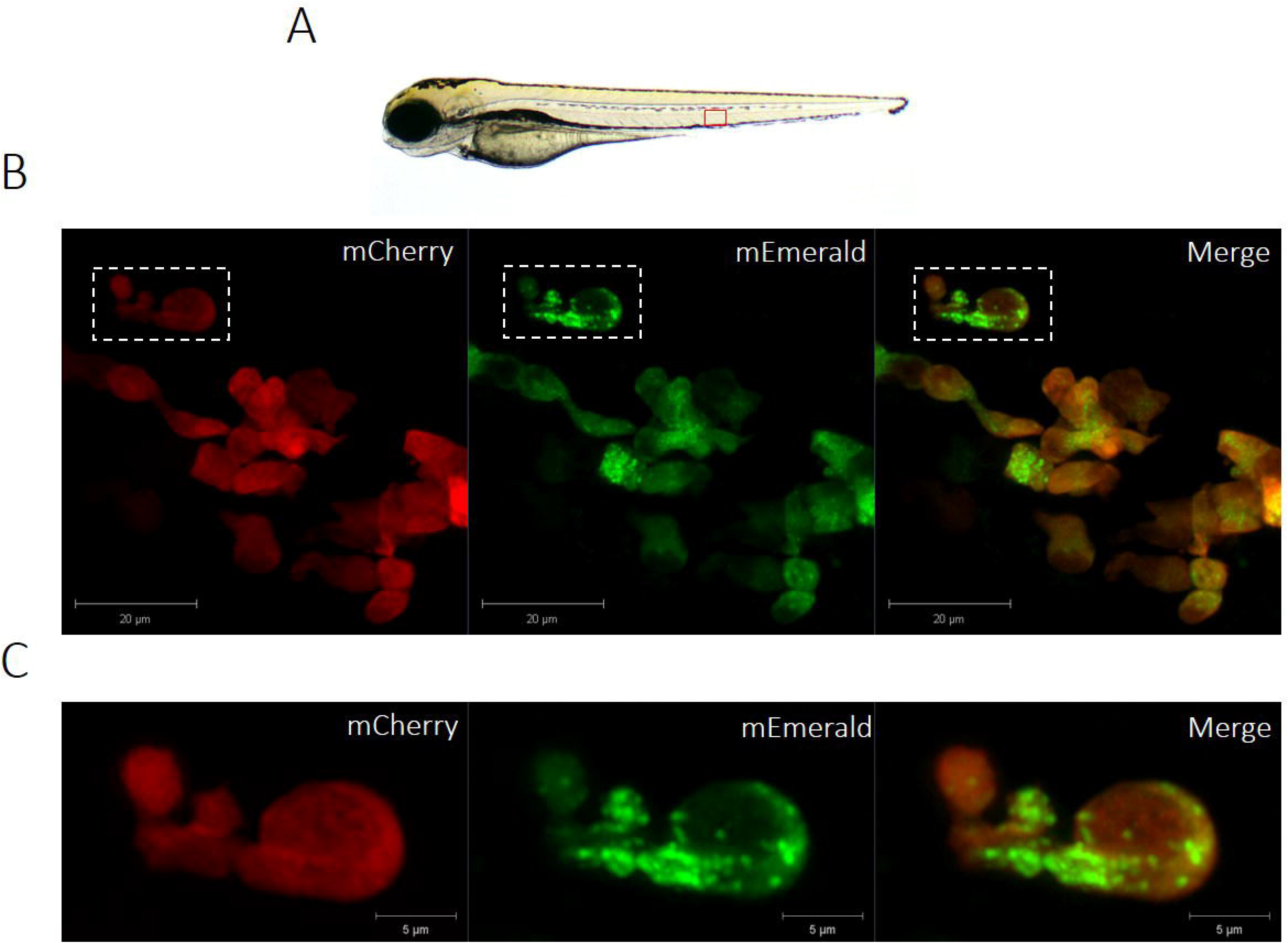
The *lyz:*MPO-mEmerald transgene labels intracellular neutrophil granules **A)** 3dpf zebrafish larva, the field of view shown in B) is outlined by the red box. **B)** An Airyscanner confocal image of neutrophils within the CHT of a double-transgenic *Tg(lyz:Hsa.MPO-mEmerald,cmlc2:EGFP)sh496; Tg(lyz:nfsB-mCherry)sh260* larva at 3dpf. **C)** An enlarged image of the neutrophil highlighted by the dashed white box in B). Scale bars **B)** 20µm and **C)** 5µm.

**Figure 4.**
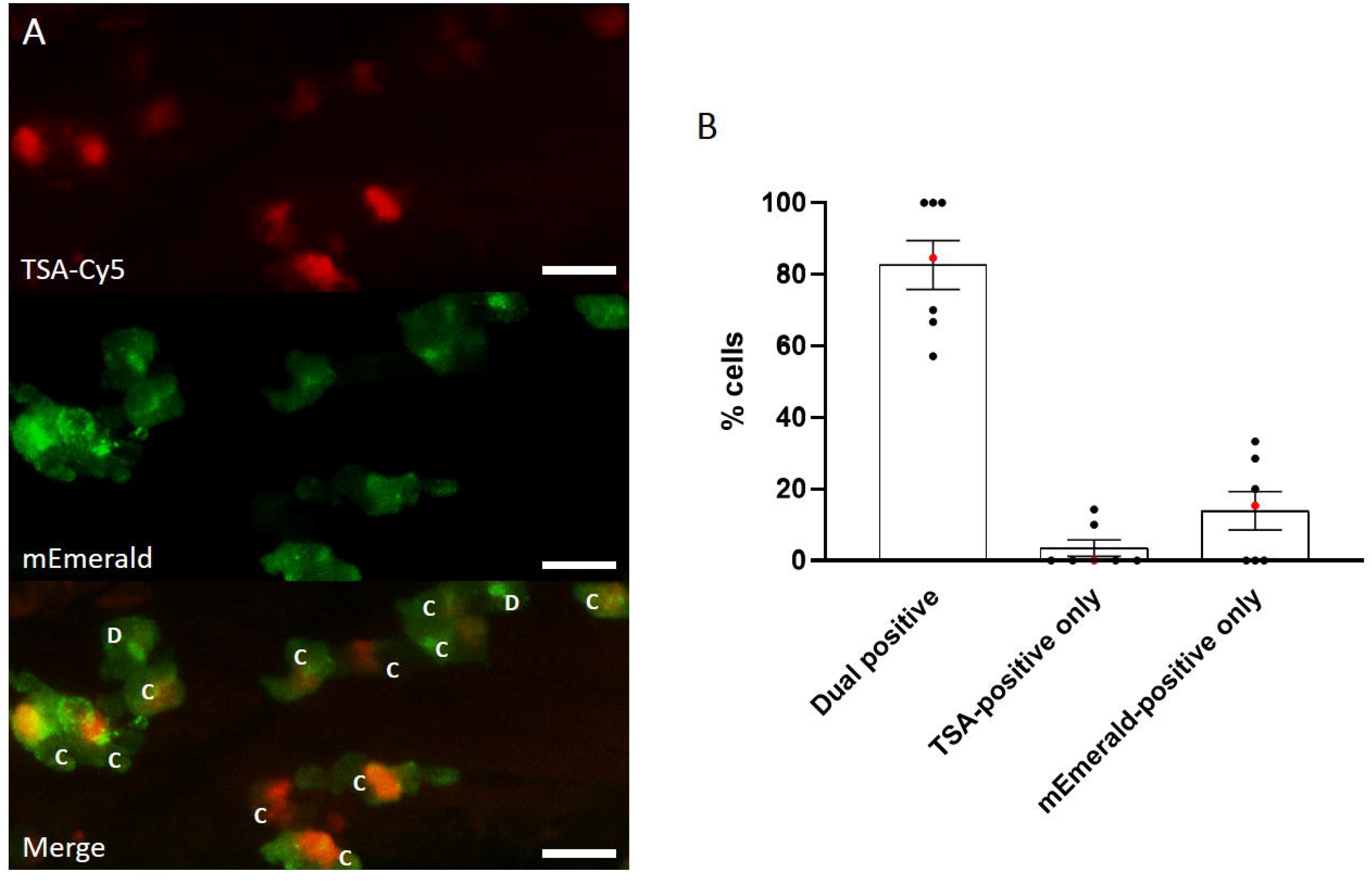
Most MPO-mEmerald-positive neutrophils are TSA-positive **A)** Confocal photomicrographs shown as maximum intensity projections of *lyz*:MPO-mEmerald larvae fixed at 3 dpf and chemically stained with TSA-Cy5. Dual-positive **(C)** and mEmerald-positive only **(D)** cells are indicated; scale bar 10µm. **B)** Quantification of dual positive, TSA-positive only and mEmerald-positive only cells observed in *lyz*:MPO-mEmerald larvae fixed at 3 dpf and chemically stained with TSA-Cy5. Red points indicate larva shown in A).

**Figure 5.**
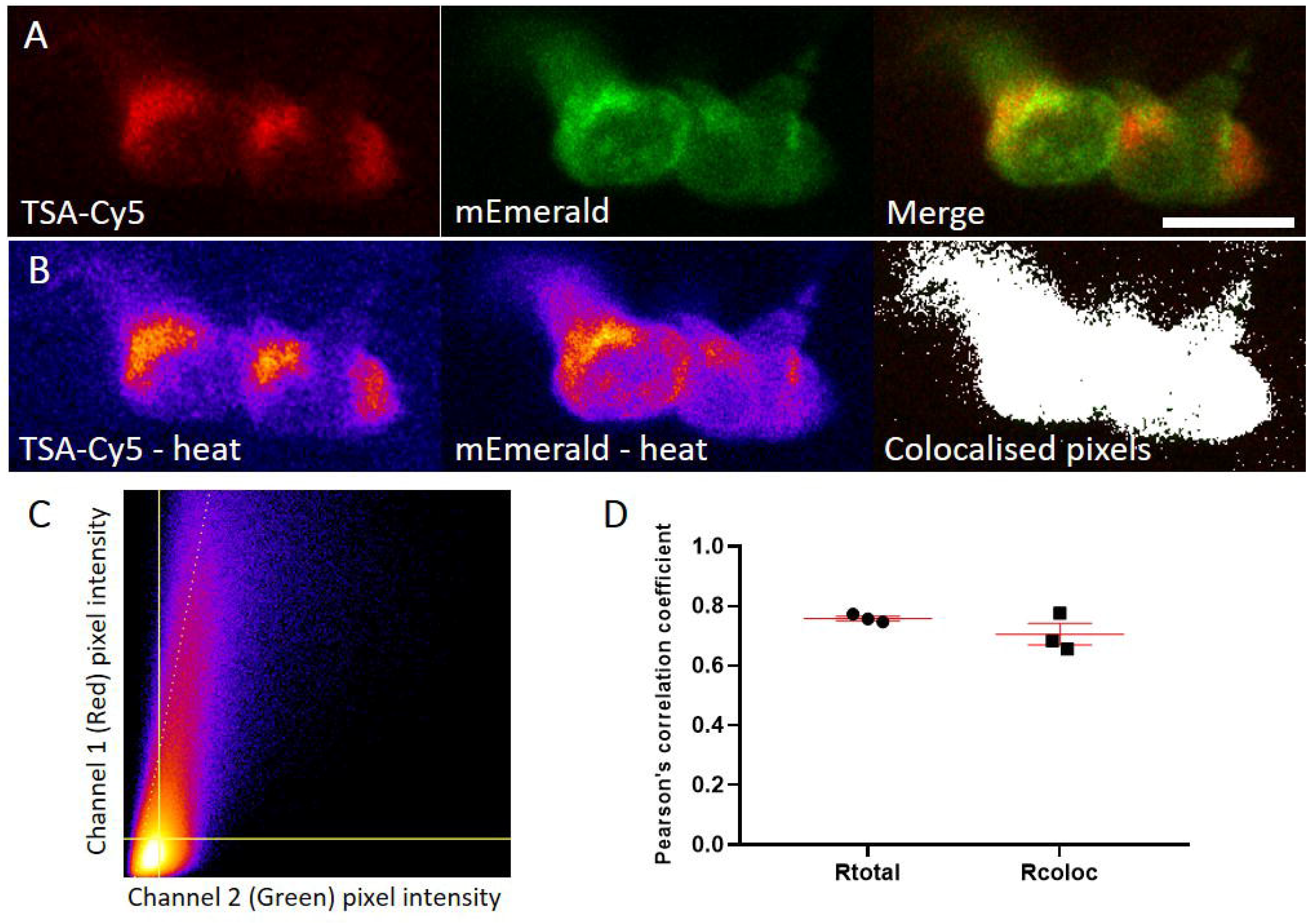
mEmerald signal colocalises with TSA signal **A)** Representative photomicrograph shown as single focal plane of *lyz*:MPO-mEmerald (Green) larvae fixed at 3 dpf and chemically stained with TSA-Cy5 (Red). Scale bar 10 µm. **B)** Pseudocoloured (heat) images of Red and Green channels and image of colocalised pixels above threshold. **C)** Scatter plot of channel 1 (Red) vs. channel 2 (Green) of the image shown in A and B. The regression line is plotted along with the threshold level for channel 1 (vertical line) and channel 2 (horizontal line). **D)** Pearson’s correlation coefficient for the entire image (Rtotal) or for the pixels above thresholds (Rcoloc) of 3 tested field of views. Mean +/- SEM are indicated in red.

### MPO-mEmerald does not disrupt neutrophil migration

In addition to its role in potentiating ROS generation in neutrophils, MPO also influences neutrophil migration to inflammatory stimuli (16). Accordingly, we sought to determine whether *Tg(lyz:Hsa.MPO-mEmerald,cmlc2:EGFP)sh496* fish exhibit disrupted neutrophil migration to inflammatory and infectious stimuli. *Tg(lyz:Hsa.MPO-mEmerald,cmlc2:EGFP)sh496* fish were crossed to *Tg(lyz:nfsB-mCherry)sh260* and at 3dpf their larvae were separated into two groups: non- humanised (*lyz:*nfsB-mCherry only) and humanised (*lyz:*MPO-mEmerald; *lyz:*nfsB-mCherry) to determine how expression of MPO-mEmerald affects these responses.

To assess neutrophil migration to inflammatory stimuli, we used a tailfin-transection model that initiates neutrophil recruitment to a vertically transected tailfin injury in zebrafish larvae (35). Non-humanised and humanised larvae were injured and neutrophil recruitment to the site of injury was imaged at 3 and 6 hours post injury (hpi) (Figure 6A). Both groups exhibited comparable migration of neutrophils to the site of injury at 3 and 6hpi (Figure 6B), suggesting that *lyz:*MPO-mEmerald does not interfere with neutrophil recruitment to sites of injury.

**Figure 6.**
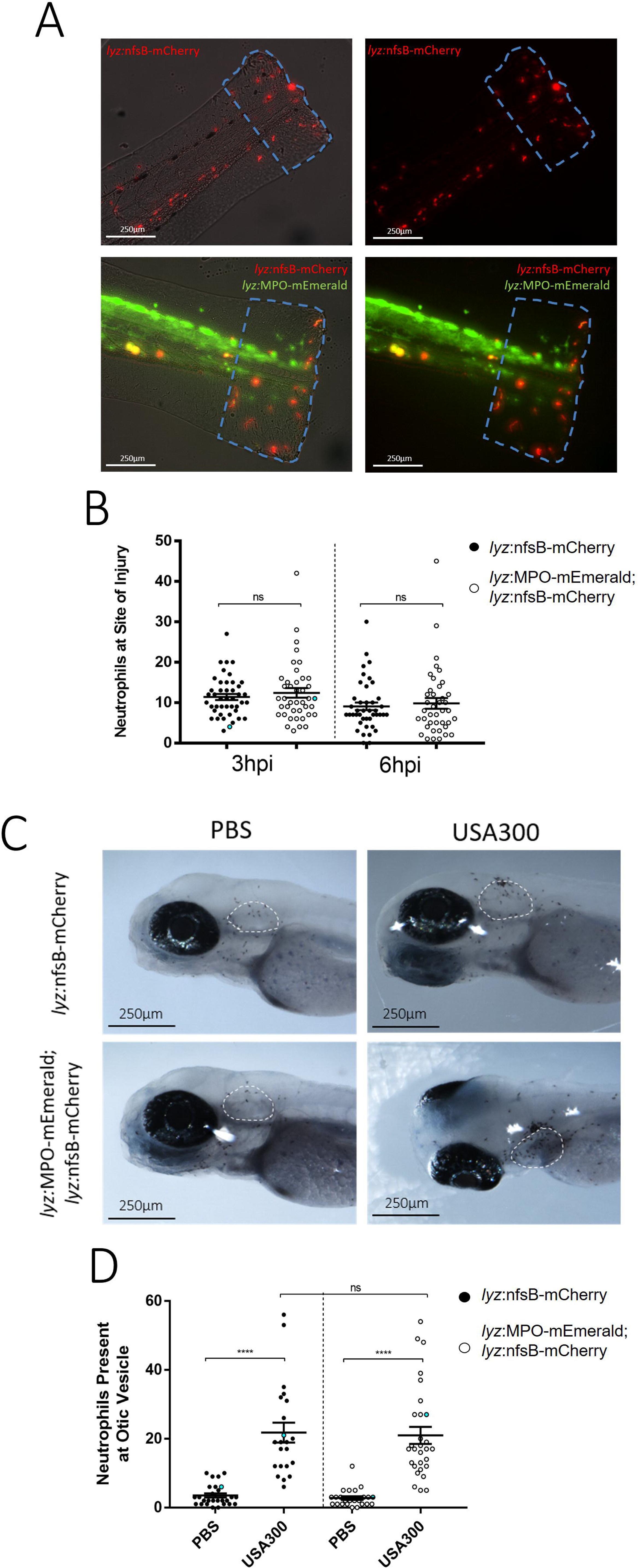
Transgene expression does not disrupt neutrophil recruitment to sites of injury or infection **A)** Non-humanised (*lyz:*nfsB-mCherry only) and humanised (*lyz:*MPO-mEmerald*; lyz:*nfsB-mCherry) 3dpf larvae with tailfins transected to induce neutrophil recruitment; dashed outline represents the area in which neutrophils were counted. Scale bar = 250µm. **B)** Neutrophils present at the site of injury at 3 and 6 hours post injury (hpi); blue points denote the representative images in A). Error bars shown are mean ± SEM (n=45 over three independent experiments); groups were analysed using an ordinary two-way ANOVA and adjusted using Bonferroni’s multiple comparisons test; ns, p>0.9999. **C)** Non-humanised and humanised larvae injected with either a PBS vehicle control or 1,400cfu *S. aureus* USA300 into the otic vesicle at 3dpf, then fixed in paraformaldehyde at 4 hours post infection (hpi) and stained with Sudan Black B to detect neutrophils; dashed white outline indicates the otic vesicle. **D)** Neutrophils present at the otic vesicle at 4hpi. Scale bars = 250µm. Error bars shown are mean ± SEM (n=25 over two independent experiments); groups were analysed using an ordinary two-way ANOVA and adjusted using Bonferroni’s multiple comparisons test. ****, p<0.0001; ns, p>0.9999.

To determine whether the neutrophil response to infection is affected by expression of *lyz:*MPO-mEmerald, we used an otic vesicle infection model to investigate neutrophil recruitment (45,46). After separating larvae into non-humanised and humanised groups, they were injected into the otic vesicle with either a PBS vehicle control or *S. aureus* USA300 at 3dpf. The larvae were then fixed in paraformaldehyde at 4 hours post infection (hpi) and stained with Sudan Black B to detect neutrophils. Injection of *S. aureus* USA300 induces robust recruitment of neutrophils to the otic vesicle, with comparable numbers recruited between non-humanised and humanised larvae (Figure 6CD). This confirms that expression of the *lyz:*MPO-mEmerald transgene does not interfere with neutrophil recruitment to sites of infection.

### Genotypic and functional identification of myeloperoxidase-null Spotless (*mpx^NL144^*) larvae

While the *lyz:*MPO-mEmerald transgene is expressed in zebrafish neutrophils in a manner that recapitulates expression in human neutrophils, it was still unknown whether MPO-mEmerald is expressed as a functional enzyme. To determine whether MPO was functional, we sought to create a zebrafish that expresses only human MPO by removing expression of the endogenous zebrafish myeloperoxidase (*mpx*) from the *Tg(lyz:Hsa.MPO-mEmerald,cmlc2:EGFP)sh496* line. This was achieved using an existing zebrafish line known as Spotless (*mpx^NL144^*), which possesses a premature stop codon in the first exon of the *mpx* gene (42). Once acquired, we aimed to cross the Spotless line to our *Tg(lyz:Hsa.MPO-mEmerald,cmlc2:EGFP)sh496* line to create a line that expresses only human MPO. Before we could create this line, it was necessary to develop a genotyping protocol that could accurately identify Spotless *mpx^NL144^* fish.

The *mpx^NL144^* allele can be identified by PCR amplification of the mutated gene from genomic DNA, followed by restriction digest of the PCR product. The restriction enzyme *Bts*CI recognises 5’ GG ATG NN 3’ sites in DNA, one of which is present within the mutated *mpx^NL144^* gene (GGA TGA) but not the wild-type *mpx^wt^* gene (GGA CGA), allowing the enzyme to determine the presence of a *mpx^NL144^* allele (Figure 7A). The PCR primers were successful in amplifying the region in the *mpx* gene from *mpx^wt^*, *mpx^NL144^* and *mpx^wt/NL144^* groups (Figure 7B), and once digested with *Bts*CI produced different DNA fragments depending on the *mpx^NL144^* allele of the fish (Figure 7C), confirming *Bts*CI digestion as an efficient means of identifying the *mpx^NL144^* allele. The accuracy of the restriction digest was confirmed further by sequencing the PCR products, confirming that the fish identified by restriction digest each have the specific basepair in the expected position (Figure 7D). After adults were genotyped, their larvae were then assessed for functional myeloperoxidase expression using the myeloperoxidase-dependent stain Sudan Black B (16) (Supplemental file 1), which verified the genotyping results (Figure 7E).

**Figure 7.**
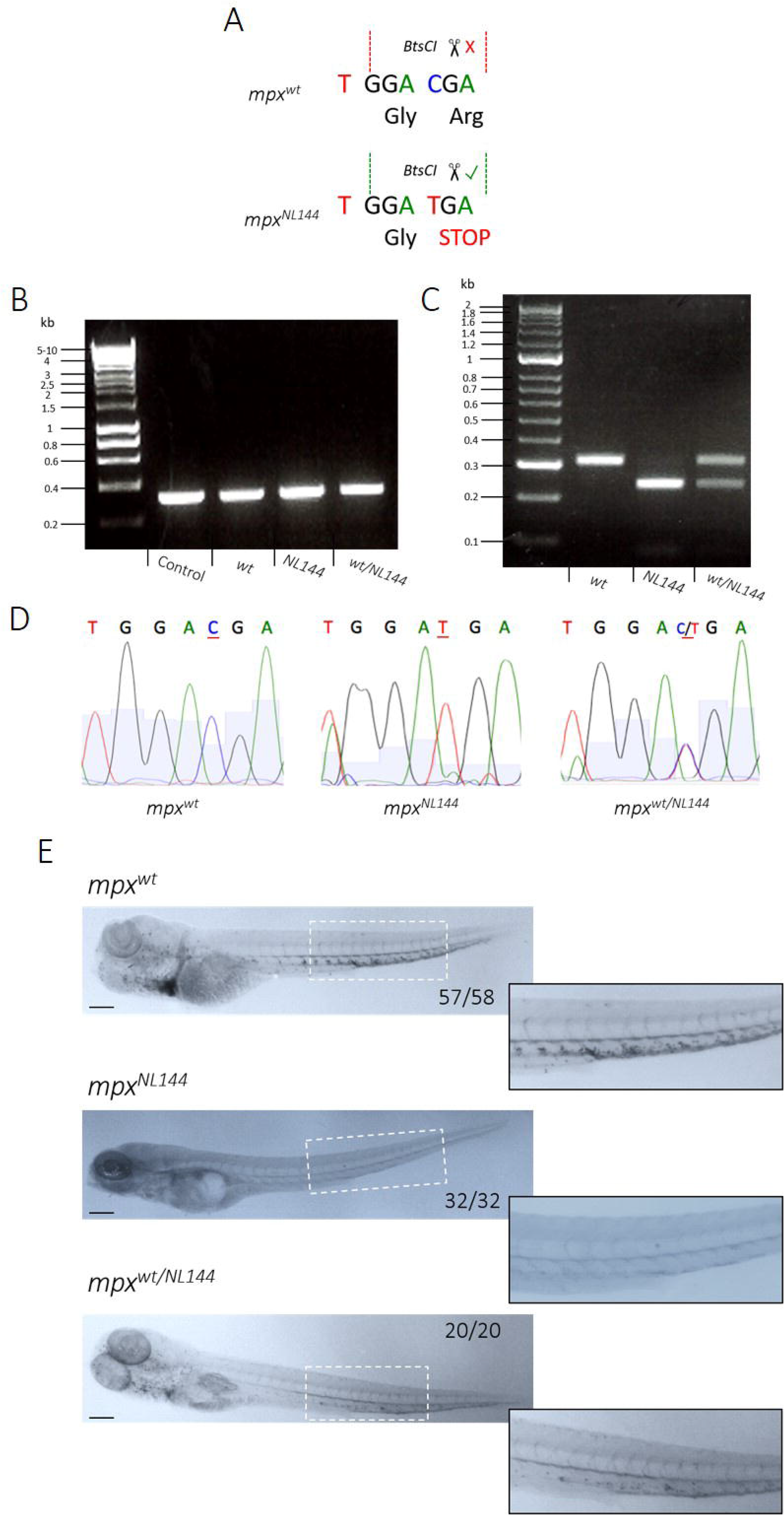
Genotyping and verifying *mpx-*null zebrafish larvae **A)** Diagram of a WT (*mpx^wt^*) and mutated (*mpx^NL144^*) gene, showing the *Bts*CI restriction site cutting only the mutated *mpx^NL144^* gene. **B)** PCR amplification of the *mpx* gene from the genomic DNA of *mpx^wt^*, *mpx^NL144^* and *mpx* fish – fragment 312bp; control DNA is a positive control from a separate genotyping experiment. *^wt/NL144^* Hyperladder 1kb. **C)** Diagnostic digest of the PCR product from *mpx^wt^*, *mpx^wt/NL144^* and *mpx*fish. Band sizes: *mpx^wt^*- 312bp, *mpx^NL144^*- 230bp, *mpx^NL144^*- 312bpand 230bp. Hyperladder 100bp plus. **D)** DNA sequencing of the PCR products to confirm the accuracy of the *Bts*CI digest. **E)** *mpx^wt^*, *mpx^NL144^* and *mpx*^wt/NL144^ larvae fixed at 4dpf and stained with Sudan Black B. Larvae with at least one functional *mpx* allele stained (57/58 *mpx^wt^*, 20/20 *mpx^wt/NL144^*) and larvae that do not produce Mpx did not stain (32/32 *mpx^NL144^*). Inset shows an enlarged view of the region indicated by the dashed white box. Scale bar = 200µm.

### MPO-mEmerald is non-functional in zebrafish neutrophils

To create zebrafish larvae expressing only human MPO, the *Tg(lyz:Hsa.MPO-mEmerald,cmlc2:EGFP)sh496* line was crossed to the Spotless line to create zebrafish that express the *lyz:*MPO-mEmerald transgene and do not produce functional endogenous *mpx^NL144^*. Once created, *Tg(lyz:Hsa.MPO-mEmerald,cmlc2:EGFP)sh496; mpx* larvae were stained with the myeloperoxidase-dependent stain Sudan Black B to determine whether this conferred staining; these larvae were compared against three sibling control groups – *mpx^NL144^*, *mpx^wt/NL144^* and *Tg(lyz:Hsa.MPO-mEmerald,cmlc2:EGFP)sh496; mpx^wt/NL144^*. All groups tested containing a functioning *mpx* allele (*mpx^wt/NL144^*, *Tg(lyz:Hsa.MPO-mEmerald,cmlc2:EGFP)sh496; mpx^wt/NL144^*) stained with Sudan Black B, indicating that the stain identifies functional endogenous myeloperoxidase (Figure 8). As expected, the negative control group did not stain (*mpx^NL144^*), but surprisingly, neither did the *Tg(lyz:Hsa.MPO-mEmerald,cmlc2:EGFP)sh496; mpx^NL144^* human MPO-only larvae, indicating that the *lyz:*MPO-mEmerald transgene does not produce a functional MPO enzyme (Figure 8).

**Figure 8.**
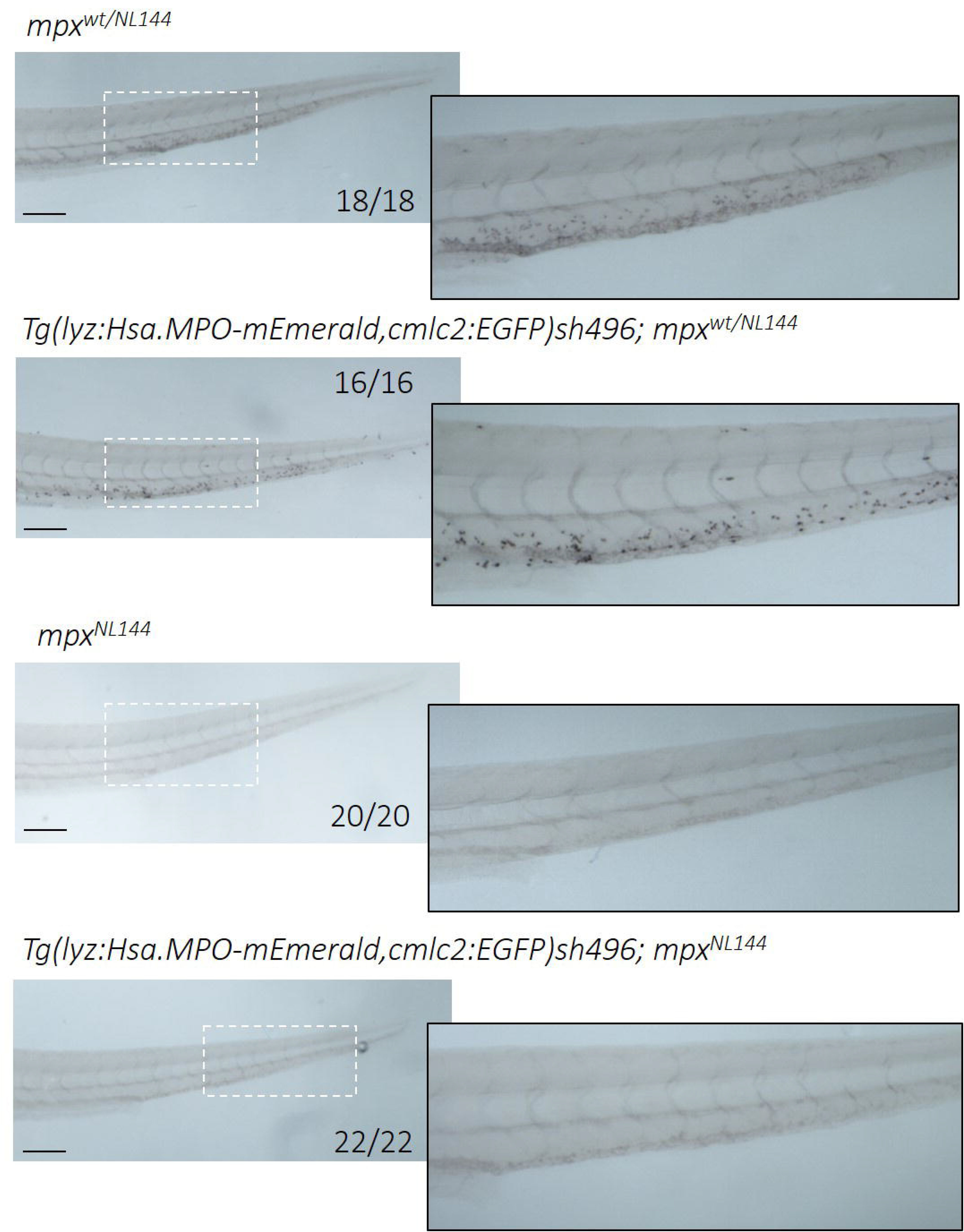
Larvae expressing only human MPO do not stain with myeloperoxidase-dependent Sudan Black B Four groups of larvae were fixed at 4dpf and stained with Sudan Black B: *mpx^wt/NL144^*, *Tg(lyz:Hsa.MPO-mEmerald,cmlc2:EGFP)sh496*; *mpx^wt/NL144^*, *mpx^NL144^* and *Tg(lyz:Hsa.MPO-mEmerald,cmlc2:EGFP)sh496*; *mpx^NL144^*. *mpx^wt/NL144^* and *Tg(lyz:Hsa.MPO-mEmerald,cmlc2:EGFP)sh496*; *mpx^wt/NL144^* stained (18/18, 16/16 respectively); *mpx^NL144^* and *Tg(lyz:Hsa.MPO-mEmerald,cmlc2:EGFP)sh496*; *mpx^NL144^* did not stain (20/20, 22/22 respectively). Dashed outline indicates the enlarged region shown adjacent. Scale bar = 200µm.

## Discussion

In this study, we created a transgenic line expressing a fluorescently-tagged human myeloperoxidase in zebrafish neutrophils. Expression in neutrophils was determined by observing expression of *lyz*:MPO-mEmerald in the fluorescent red neutrophil line, *Tg(lyz:nfsB-mCherry)sh260*, which expresses mCherry in the cytoplasm of zebrafish neutrophils. Both transgenes were expressed within the same cells (Figure 1, 2), and TSA staining showed that the majority of MPO-mEmerald cells produced active peroxidase that colocalises with MPO-mEmerald signal (Figure 4) confirming that the *lyz:*MPO-mEmerald transgene labels neutrophils. However, as MPO localises with the primary granules of neutrophils, it was essential that the fluorescent signal observed in the *lyz:*MPO-mEmerald line should differ from the cytoplasmic signal observed in the *Tg(lyz:nfsB-mCherry)sh260* line. This was observed in several instances; in double transgenic neutrophils, distinct areas of the cell remain unlabelled with mEmerald (Figures 1, 3) suggesting that MPO is translated and trafficked to a subcellular location that is distinct from the cytoplasm. This observation is also evident in Airyscan confocal imaging (Figure 3C), where a large unlabelled area of a double-transgenic neutrophil is visible in the mEmerald channel. This is likely to be a region of the cell that is inaccessible to the primary granules, for example the nucleus, and could be verified using a fluorescent nuclear probe.

In addition to the *lyz:*MPO-mEmerald and *lyz:*nfsB-mCherry signals being distinct, double-transgenic neutrophils contain small intracellular foci of mEmerald signal (Figure 3), suggesting that MPO-mEmerald might be targeted to the primary granules. Using high-speed imaging, we found that these foci are highly dynamic, resembling the Brownian motion exhibited by neutrophil granules (Supplemental movie 1). Despite these observations, we found that an average of 80% of cells expressing MPO-mEmerald stain with peroxidase-sensitive TSA, and 70% of mEmerald signal colocalises with TSA signal (Figure 5). It is important to note that not all labelled MPO would localise with active peroxidase, as immature MPO present in the endoplasmic reticulum and trans-golgi network prior to dimerisation would also be visible in MPO-mEmerald fish (47). Additionally, it is likely that neutrophils at different stages of development would contain different levels of functional MPO, which may explain incomplete staining with TSA.

In addition to the role of MPO in antibacterial defence, it is also an important enzyme regulating the migration of neutrophils to sites of infection and inflammation, primarily by mediating H_2_O_2_ flux (16). Using a combination of approaches for studying neutrophil migration, we found that expression of the *lyz*:MPO-mEmerald transgene does not interfere with neutrophil recruitment to sites of infection and inflammation (Figure 6). Currently, there are no existing tools for visualising neutrophil granules *in vivo*, and measuring MPO is limited to cytochemical and cytometry-based approaches (36). Therefore, the *Tg(lyz:Hsa.MPO-mEmerald,cmlc2:EGFP)sh496* transgenic line may be used to study granule dynamics *in vivo* without disrupting neutrophil function.

In order to determine whether MPO-mEmerald is produced as a functional enzyme, it was necessary to produce zebrafish that do not express endogenous zebrafish myeloperoxidase. We describe here a genotyping protocol that can be used to identify Spotless (*mpx^NL144^*) fish, an *mpx*-null mutant line created in a separate study (42). This was accomplished by amplifying a region present in the first exon of the *mpx* gene, followed by restriction digest with *Bts*CI (Figure 7BCD); this was then functionally verified using the myeloperoxidase-dependent stain Sudan Black B (16) (Figure 7E). We believe this to be a useful and robust method for identifying Spotless fish, and may be useful in future studies.

Once a robust method for identifying Spotless fish was established, the Spotless line was crossed to the *Tg(lyz:Hsa.MPO-mEmerald,cmlc2:EGFP)sh496* line to generate a line that expresses human MPO, and does not express zebrafish Mpx (known as *Tg(lyz:Hsa.MPO-mEmerald,cmlc2:EGFP)sh496; mpx^NL144^*). By comparing staining with Sudan Black B with sibling controls, we found that MPO-mEmerald is not produced as a functional enzyme, as *lyz:*MPO-mEmerald expression does not complement staining in the *mpx^NL144^* background. It is unclear why the *Tg(lyz:Hsa.MPO-mEmerald,cmlc2:EGFP)sh496* line does not produce functional MPO, however it is important to note that MPO is a complex glycoprotein enzyme that undergoes numerous tightly regulated post-translational modifications. Before mature MPO is produced, the peptide associates with calreticulin and calnexin in the endoplasmic reticulum before undergoing a series of proteolytic events leading to insertion of a haem group and dimerisation of the enzyme, followed by glycosylation and ending with granule targeting (47). The importance of each step in producing a functional enzyme is unclear, however studies of myeloperoxidase-deficient individuals suggest that targeting to the primary granules universally correlates with functional MPO (23,48–50), and *in vitro* studies show that dimerisation is not required for enzyme function (15,51,52). Additionally, the discrepancy is unlikely to lie with calnexin and calreticulin, as they possess roughly 70% amino acid identity with the human chaperones, and are important during development of the zebrafish lateral line (53). Differences at any other stages may lead to incomplete MPO maturation and function in the zebrafish and consequently, the *Tg(lyz:Hsa.MPO-mEmerald,cmlc2:EGFP)sh496* line may also be useful in investigating how MPO is processed and targeted to the granules during development.

## Conclusion

We have generated a transgenic zebrafish line expressing fluorescently labelled human MPO within its neutrophils. The enzyme is non-functional and does not interfere with neutrophil recruitment to sites of infection or inflammation, suggesting that it may be used to study granule dynamics *in vivo* without disrupting neutrophil behaviour. Additionally, the *Tg(lyz:Hsa.MPO-mEmerald,cmlc2:EGFP)sh496* line may be used to investigate processing and targeting of MPO during development, which is currently uncharacterised *in vivo*. Lastly, we provide a protocol for genotyping endogenous myeloperoxidase-null Spotless (*mpx^NL144^*) fish, which will prove useful in future studies investigating myeloperoxidase in the zebrafish.

## Supporting information

Raw data: Figure 1

Raw data: Figure 2

Raw data: Figure 3

Raw data: Figure 4 & 5

Raw data: Figure 6

Raw data: Figure 7 & 8

## Acknowledgements

K.D.B performed experiments with assistance from T.K.P, M.v.G, N.W.M.d.J, and J.K. S.A.R, J.A.G.v.S and S.J.F conceived the study and designed experiments. T.K.P performed TSA-staining and colocalisation experiments. K.D.B and S.A.R wrote the manuscript with significant input from all authors. Thank you to Annemarie Meijer for providing the Spotless (*mpx^NL144^*) line, to the Bateson Centre aquarium staff for all their help, and to the Wolfson Light Microscopy Facility.

---

Supplemental movie 1. MPO-mEmerald signal is highly dynamic An airyscanner confocal timelapse of a double-transgenic *Tg(lyz:Hsa.MPO-mEmerald,cmlc2:EGFP)sh496; Tg(lyz:nfsB-mCherry)sh260* larva. The timelapse shows numerous neutrophils in the caudal haematopoietic tissue of a 3dpf larva.

Supplemental file 1. Sudan Black B Staining Protocol

