## Supplemental file 1 for "A transgenic zebrafish line for in vivo visualisation of neutrophil myeloperoxidase"

### Sudan Black Staining Protocol (7/12/15)

Reagents;

- Sudan Black B Staining Solution (Sigma)
- 4% Paraformaldehyde (Pfa) – needs to defrost for a couple of hours, do not use if defrosted over a week before.
- Phosphate Buffered Saline (PBS)
- 70% Ethanol
- PBS-Tween (0.1% Tween) (PBST)
- 30% Glycerol (0.1% Tween):
  - To make 50ml = 15ml Glycerol, 50ul Tween, to 50ml ddH_2_O
- 80% Glycerol (0.1% Tween):
  - To make 50ml = 40ml Glycerol, 50ul Tween, to 50ml ddH_2_O
- Bleaching Solution (make without H_2_O_2_ then add H_2_O_2_ last):
  - 0.5x SSC, 5% Formamide, 10% H_2_O_2_ (of 30% H_2_O_2_ max stock)

Method

1. Grow embryos for 3 days or more: (optional) add ptu to the media (E3) to remove pigmentation, alternatively bleach embryos at the end.
2. After anaesthetising embryos using 1/20 Tricaine (roughly 1ml to a petri dish), place a maximum of 20 embryos in a 1.5ml eppendorf tube and wait until embryos have sunk to the bottom.
3. Draw off as much media as possible without disturbing embryos, and then add 1ml of room-temperature Pfa.
4. Leave the eppendorf on its side and allow the embryos to fix for at least an hour at room temperature (longer fixation is fine, can go over the weekend in fix for example; leave triton or tween out of the fix solution).
5. Rinse 3x 5min in PBS by drawing off Pfa, adding 1ml of PBS and leaving on its side for 5 minutes each time.
6. Draw off PBS and add 500ul Sudan Black Staining solution for 20 minutes.
7. Carefully draw off staining solution as embryos will not be visible through the opaque solution; it’s not important that all the solution is drawn off, as embryos will be visible from the first wash. Dispose of all Sudan Black waste in a 50ml falcon throughout the staining procedure.
8. De-stain using 70% Ethanol. 4x fast rinses adding 1ml of Ethanol and drawing off. On fourth wash (or as long as it takes for clarity to return), leave on its side for 1 hour to soak.
9. Re-hydrate embryos by adding PBST. Leave roughly 300ul of 70% Ethanol and add 300ul PBST to the Ethanol, leave for 5 minutes, draw off and wash again by adding 1ml PBST for five minutes.
10. Pigmentation can be removed by adding bleaching solution to the embryos and incubating for an hour at room temperature. Then wash away H_2_O_2_ 4x 5mins in PBS. If storing long-term, keep in glycerol, for short-term storage (no more than a week) Pfa will suffice.
11. For added clarity you can clear the embryos in glycerol series: add 30% Glycerol 0.1% Tween, then draw off and add 80% Glycerol 0.1% Tween. Then place in 24-well plate.
