## Supplementary figures and images for "A transgenic zebrafish line for in vivo visualisation of neutrophil myeloperoxidase"

### 10x 1of5 BF.tif

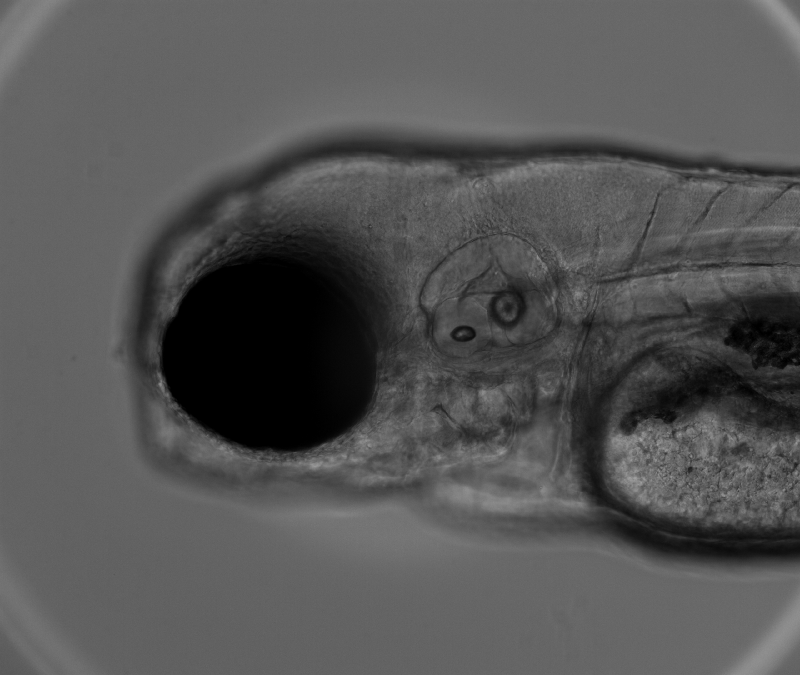

### 10x 1of5 GFP.tif

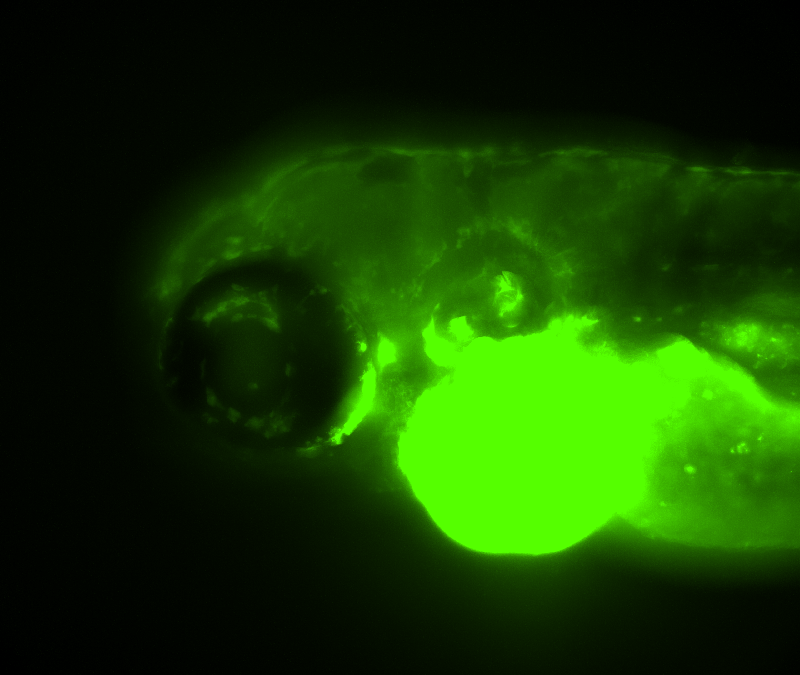

### 10x 1of5 mCherry.tif

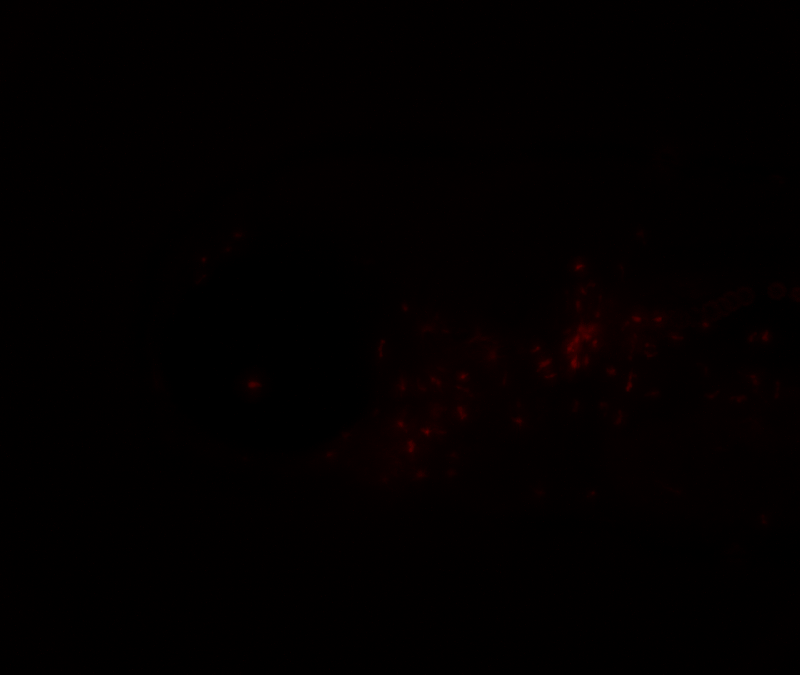

### 10x 1of5 merge.tif

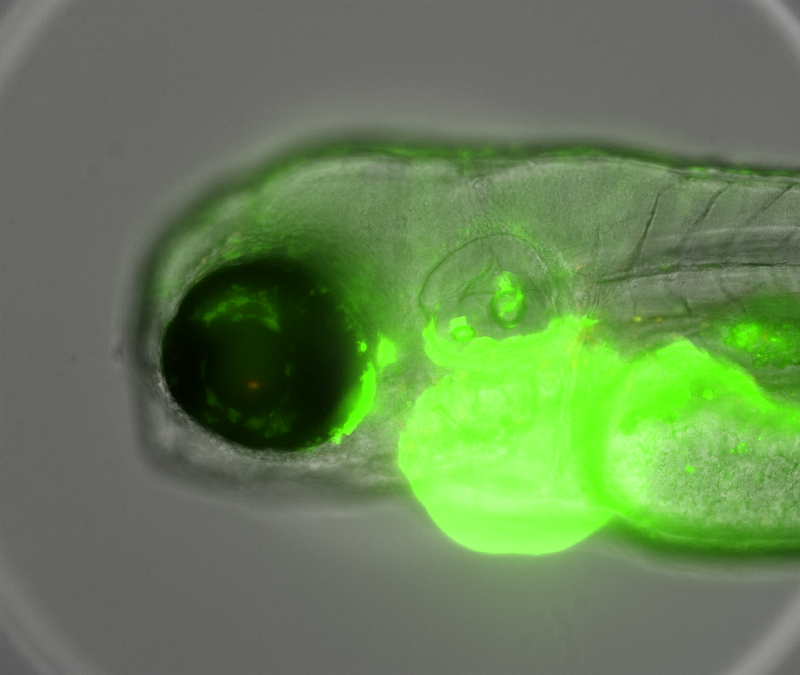

### 10x 2of5 BF.tif

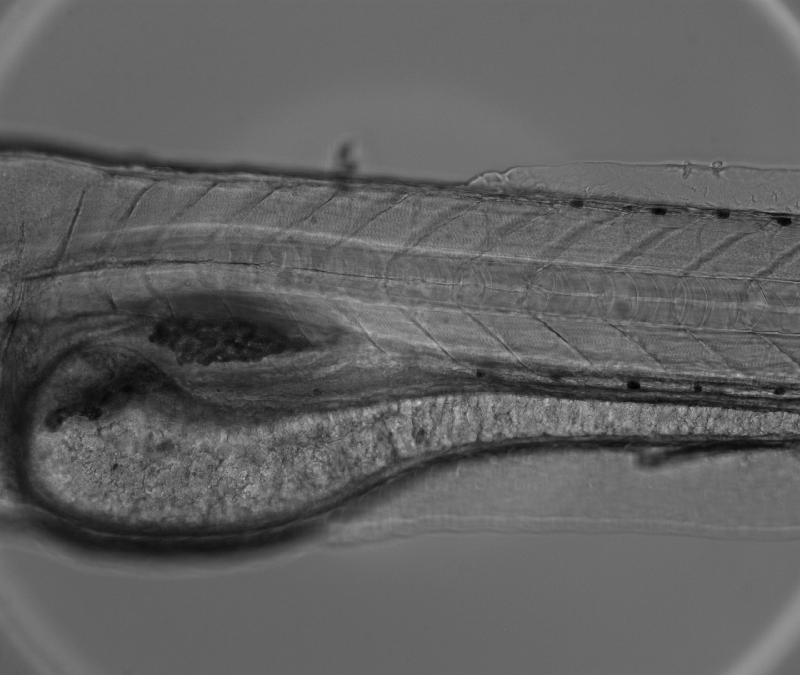

### 10x 2of5 GFP.tif

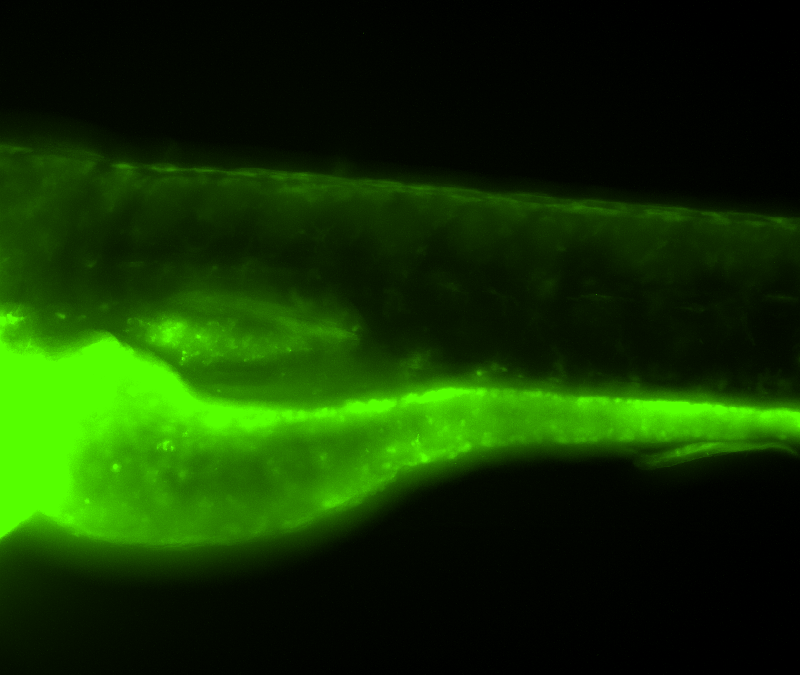

### 10x 2of5 mCherry.tif

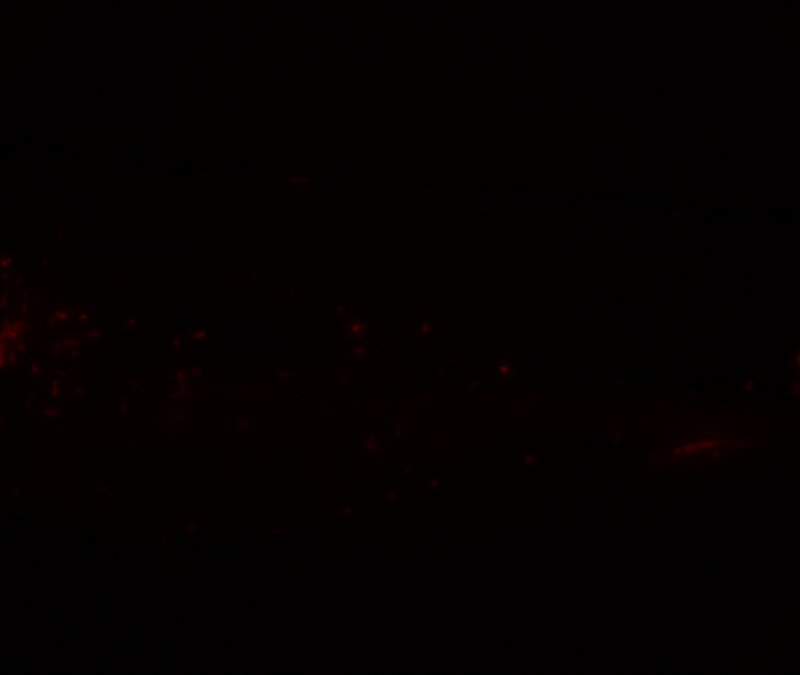

### 10x 2of5 merge.tif

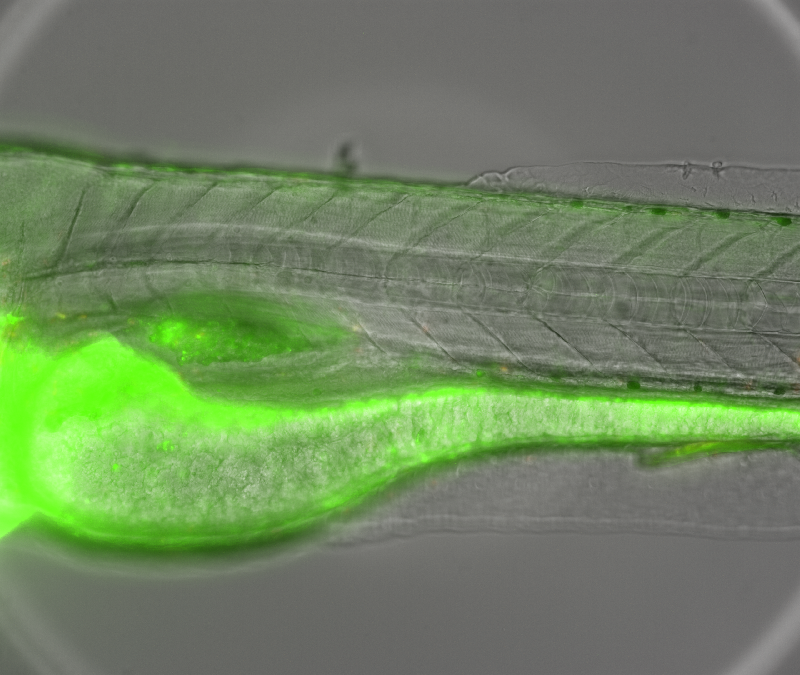

### 10x 3of5 BF.tif

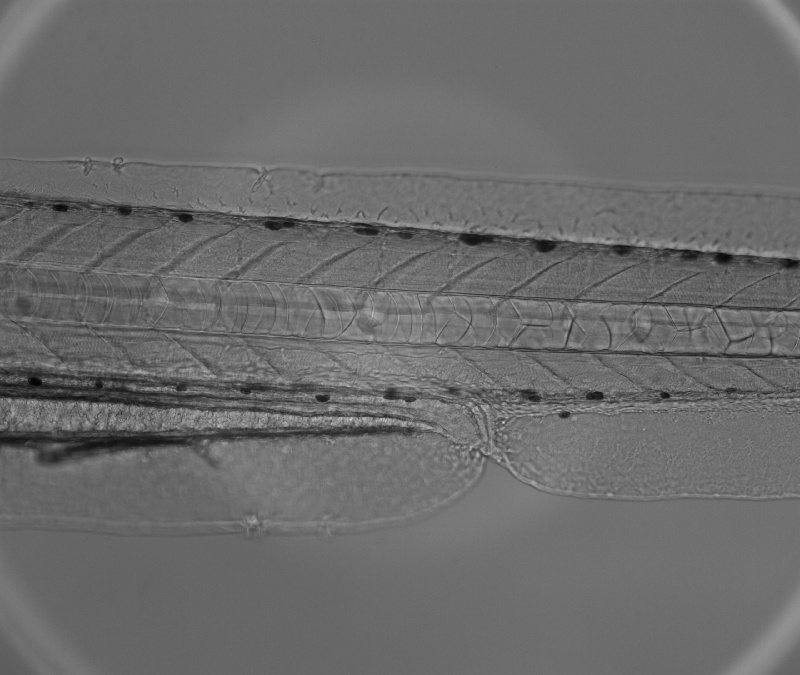

### 10x 3of5 GFP.tif

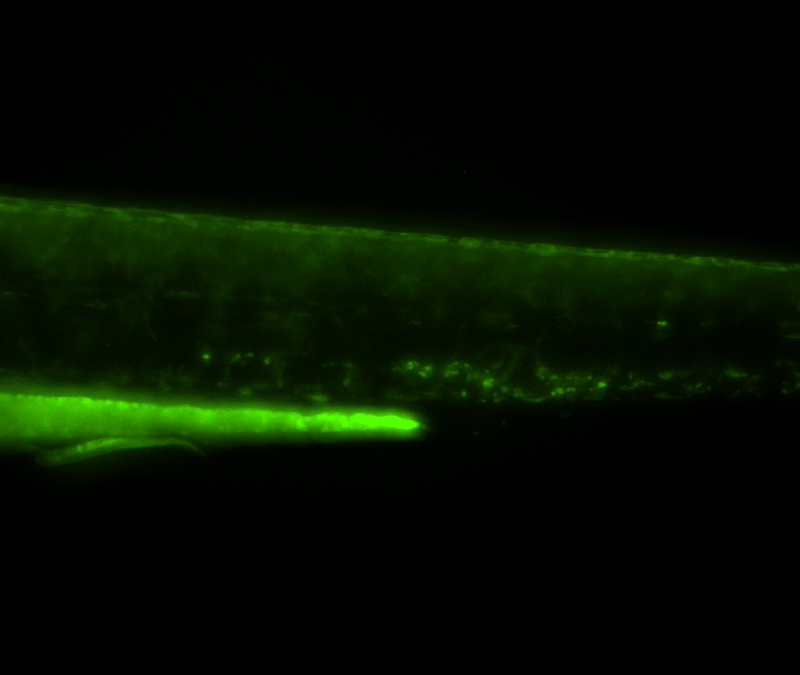

### 10x 3of5 mCherry.tif

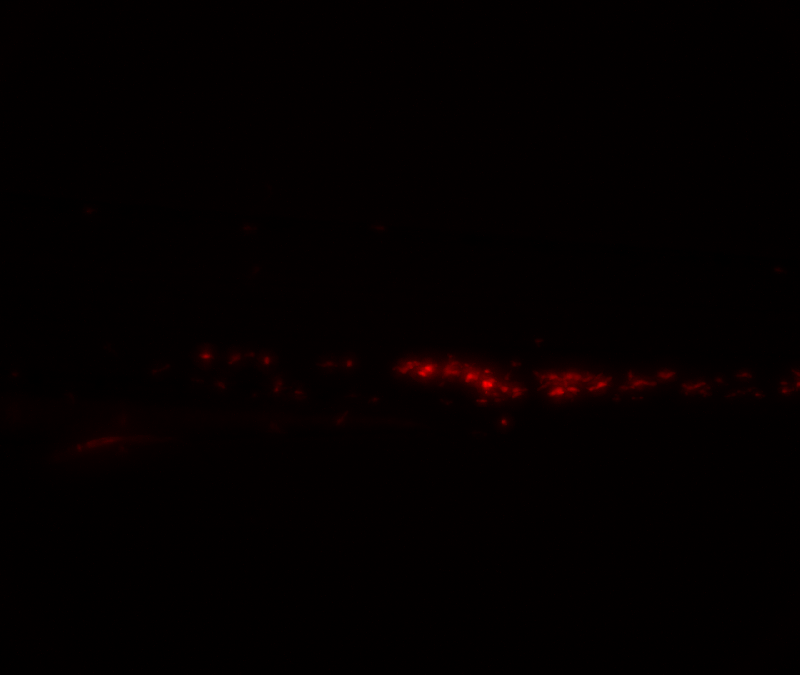

### 10x 3of5 merge.tif

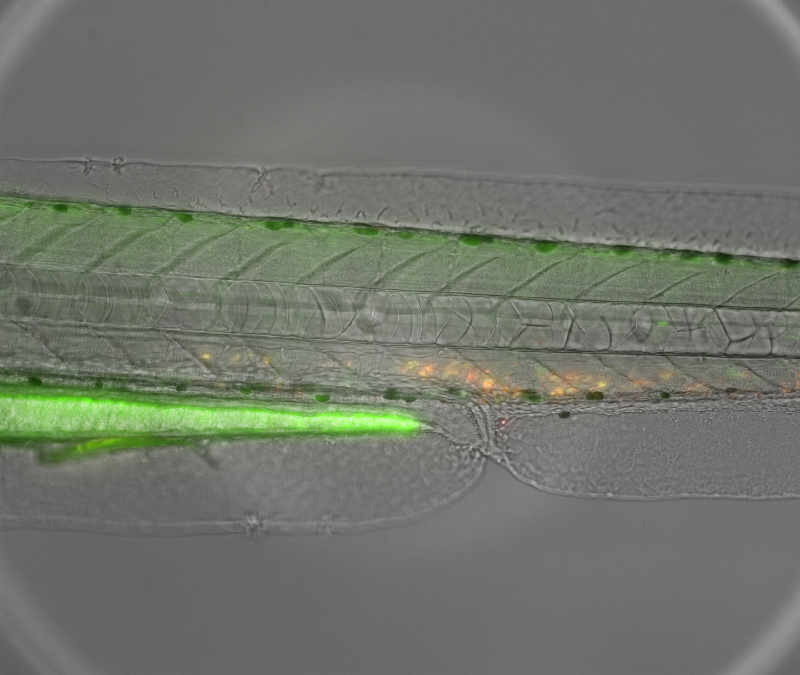

### 10x 4of5 BF.tif

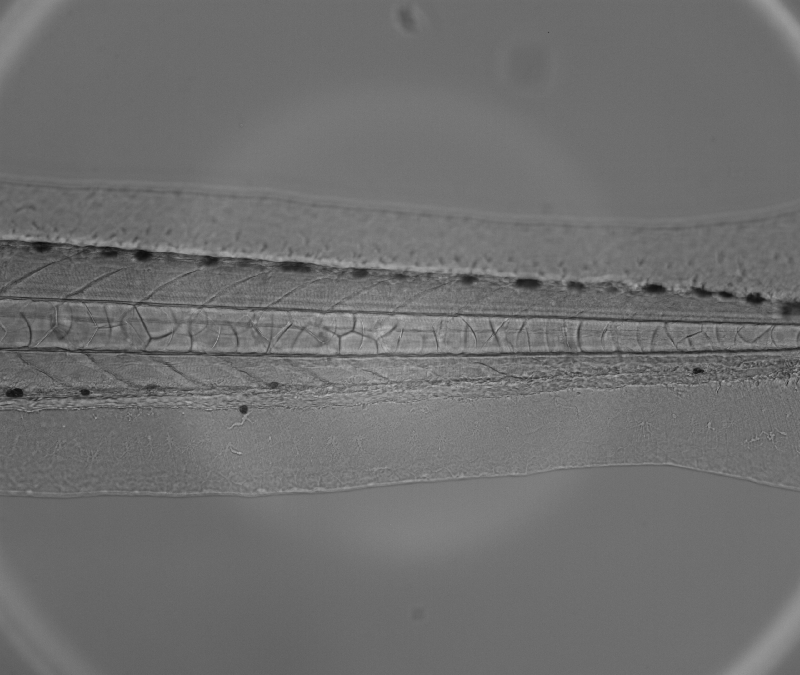

### 10x 4of5 GFP.tif

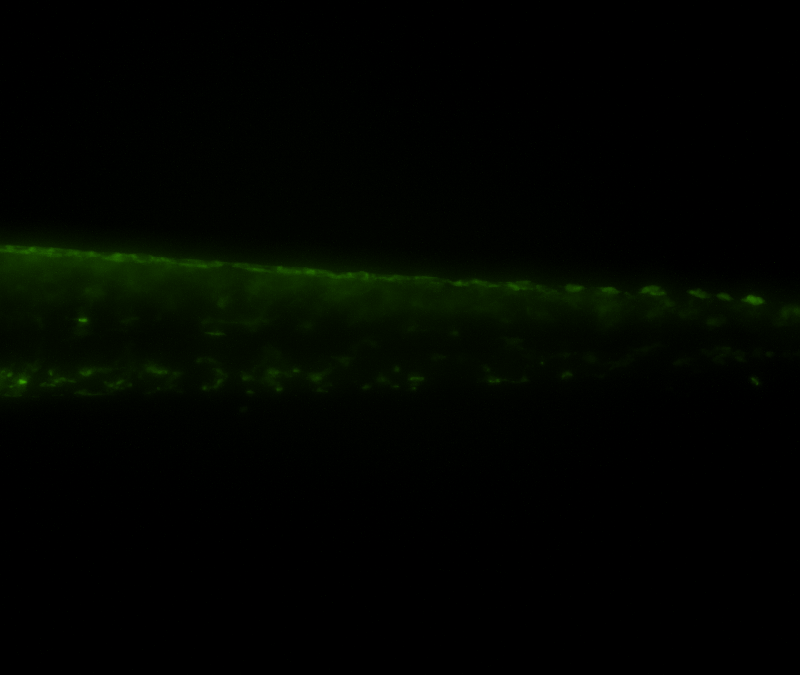

### 10x 4of5 mCherry.tif

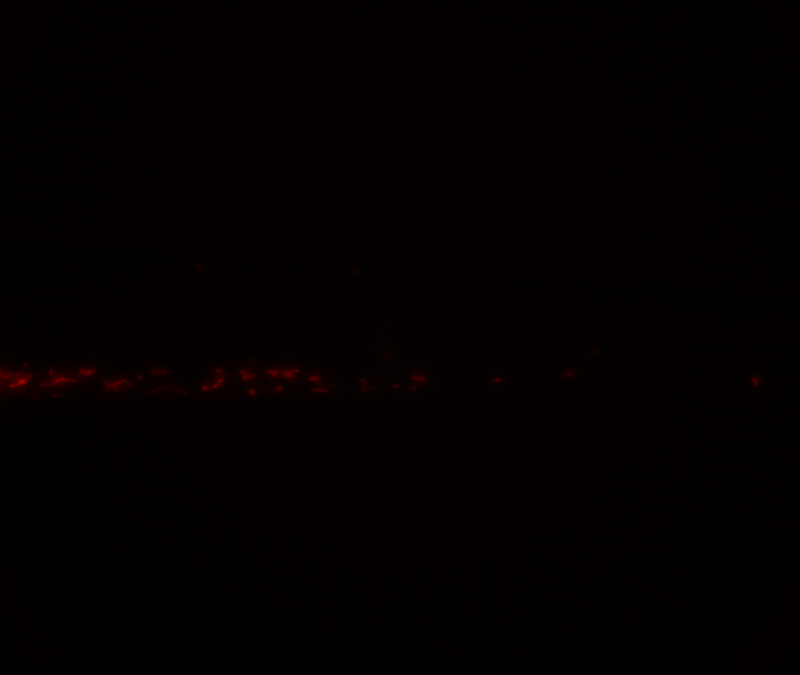

### 10x 4of5 merge.tif

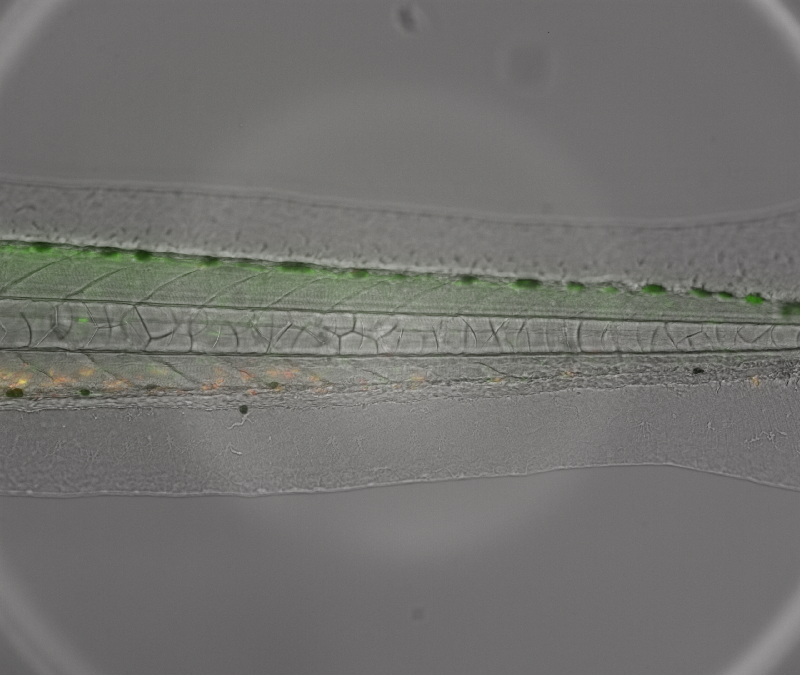

### 10x 5of5 BF.tif

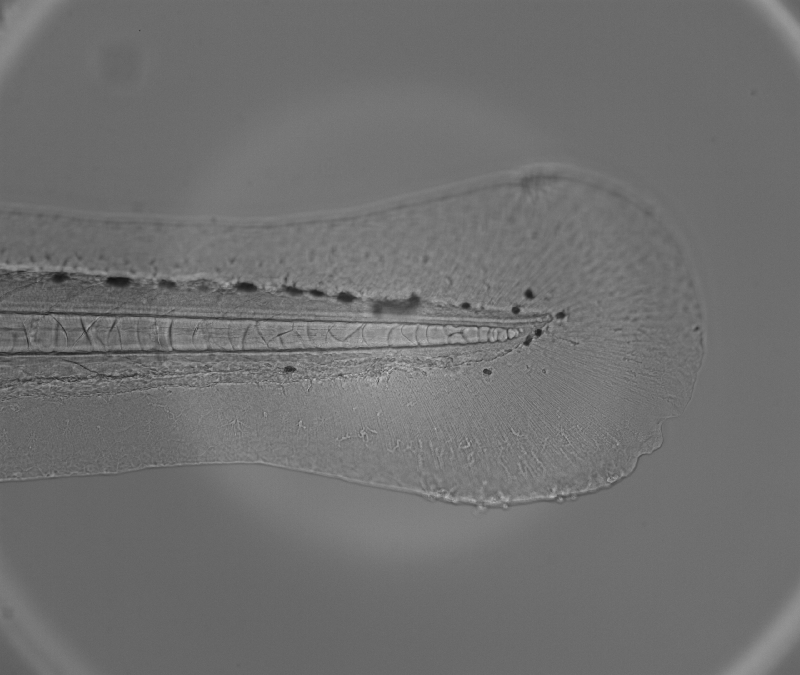

### 10x 5of5 GFP.tif

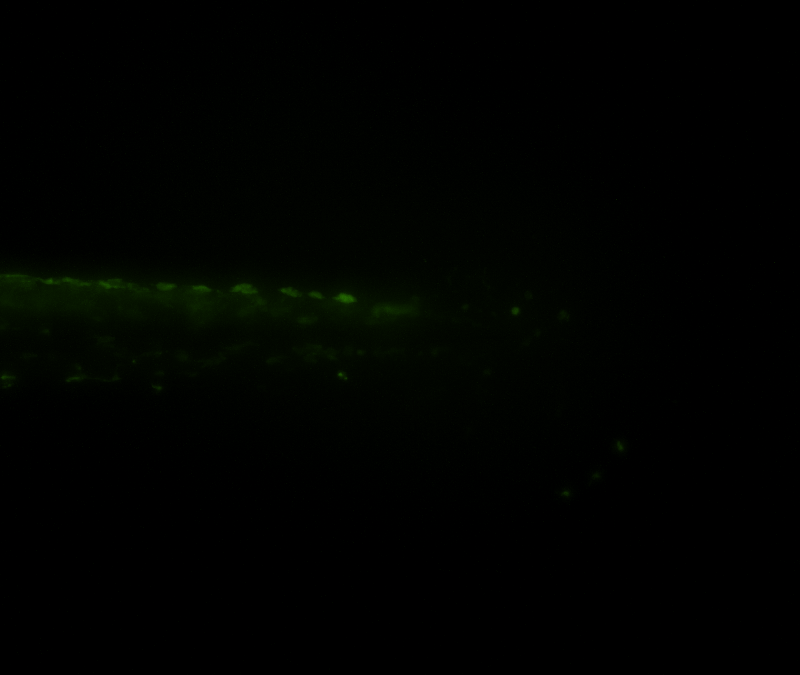

### 10x 5of5 mCherry.tif

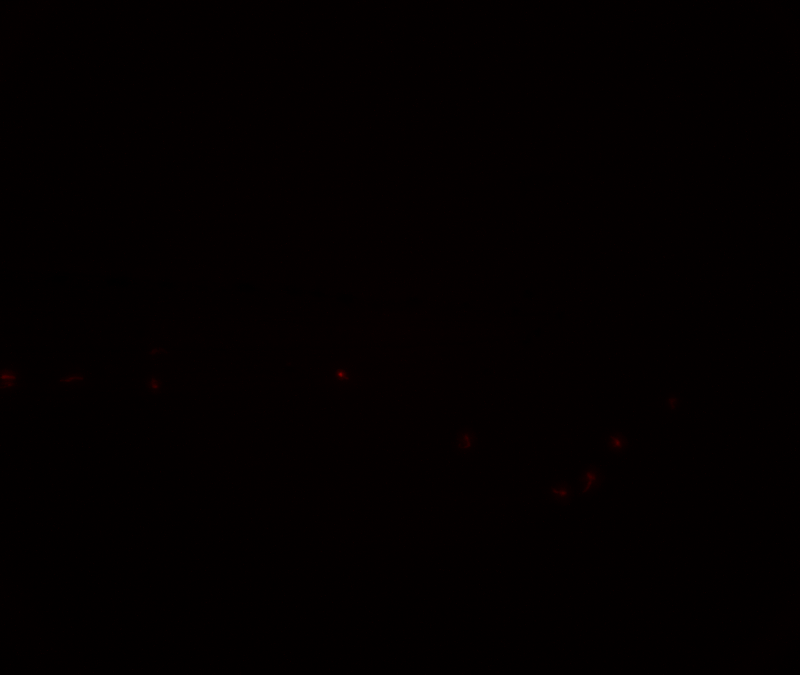

### 10x 5of5 merge.tif

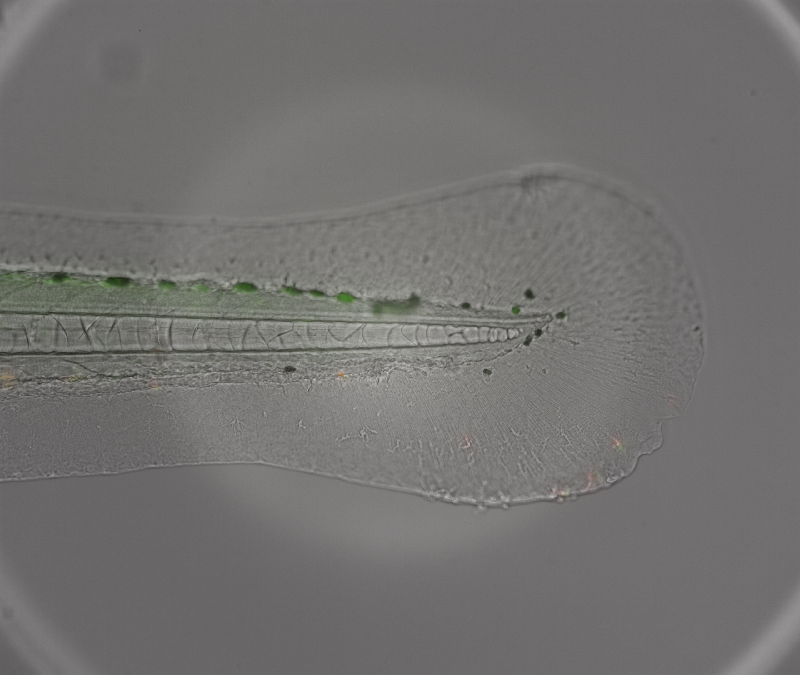

### 10x BF.tif

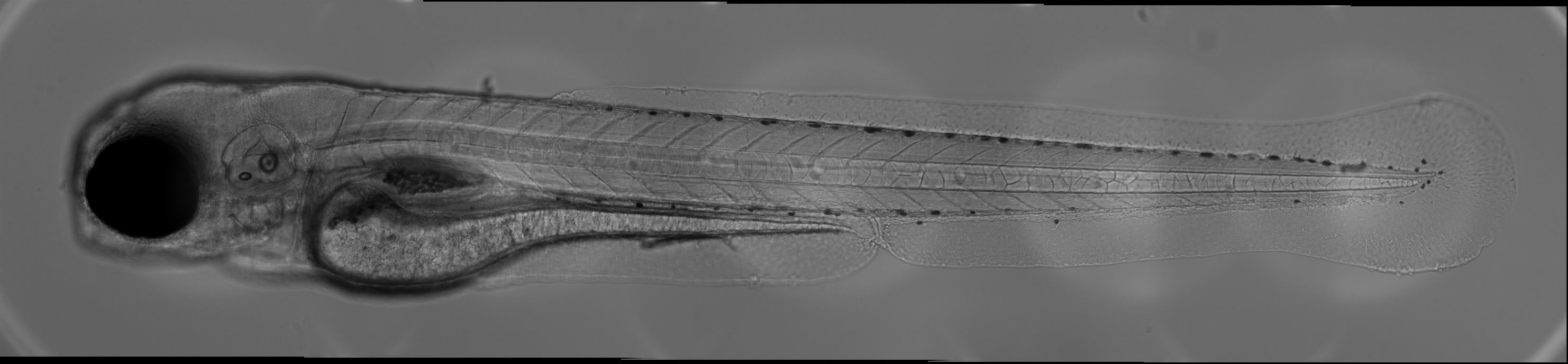

### 10x GFP.tif

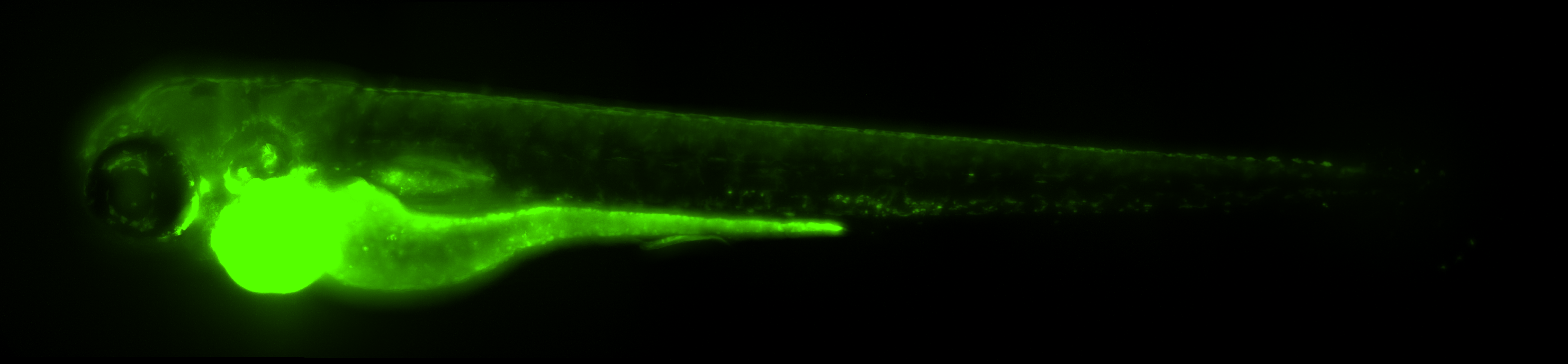

### 10x mCherry.tif

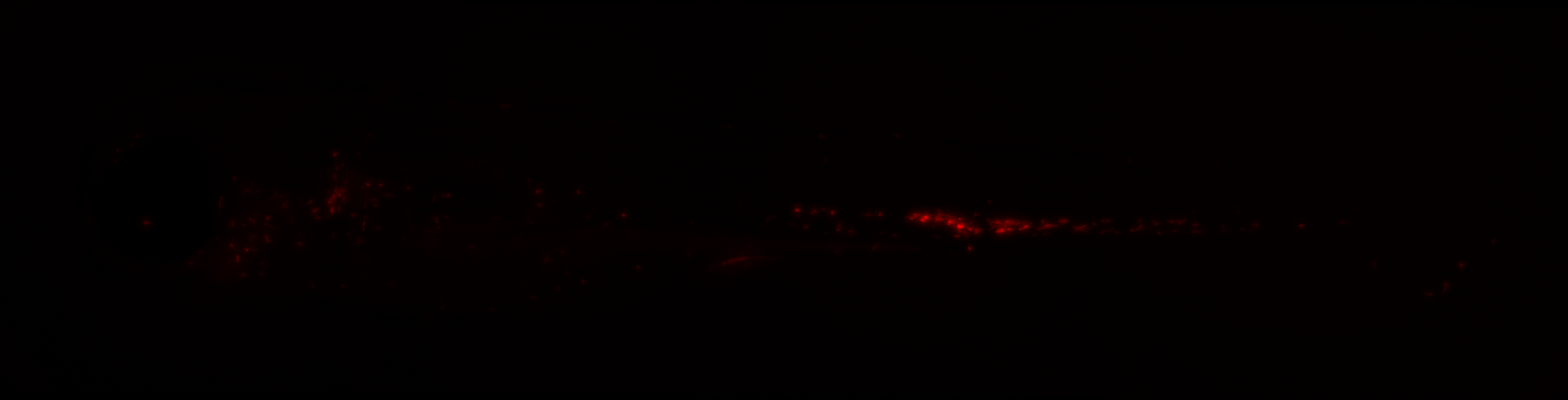

### 10x merge.tif

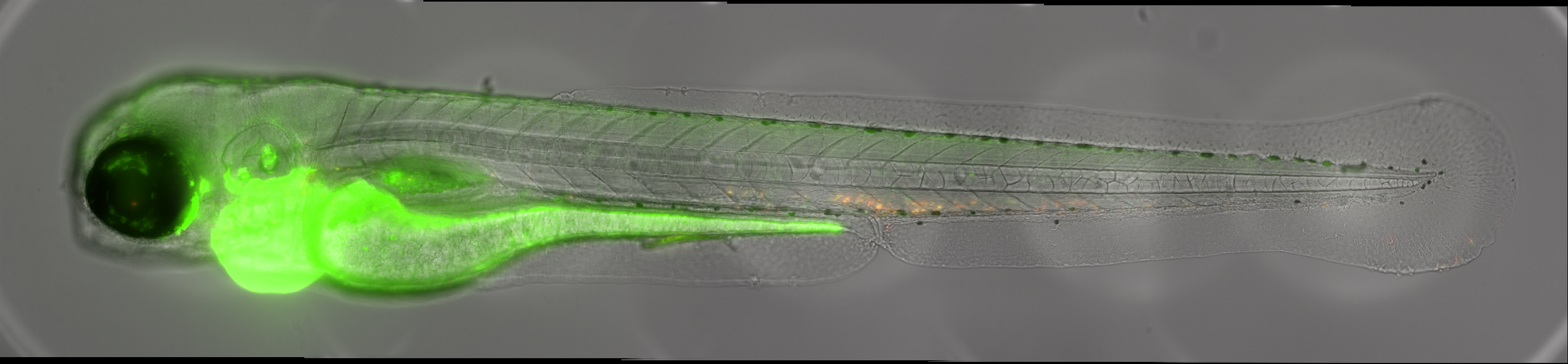

### greyscale GFP signal.jpg

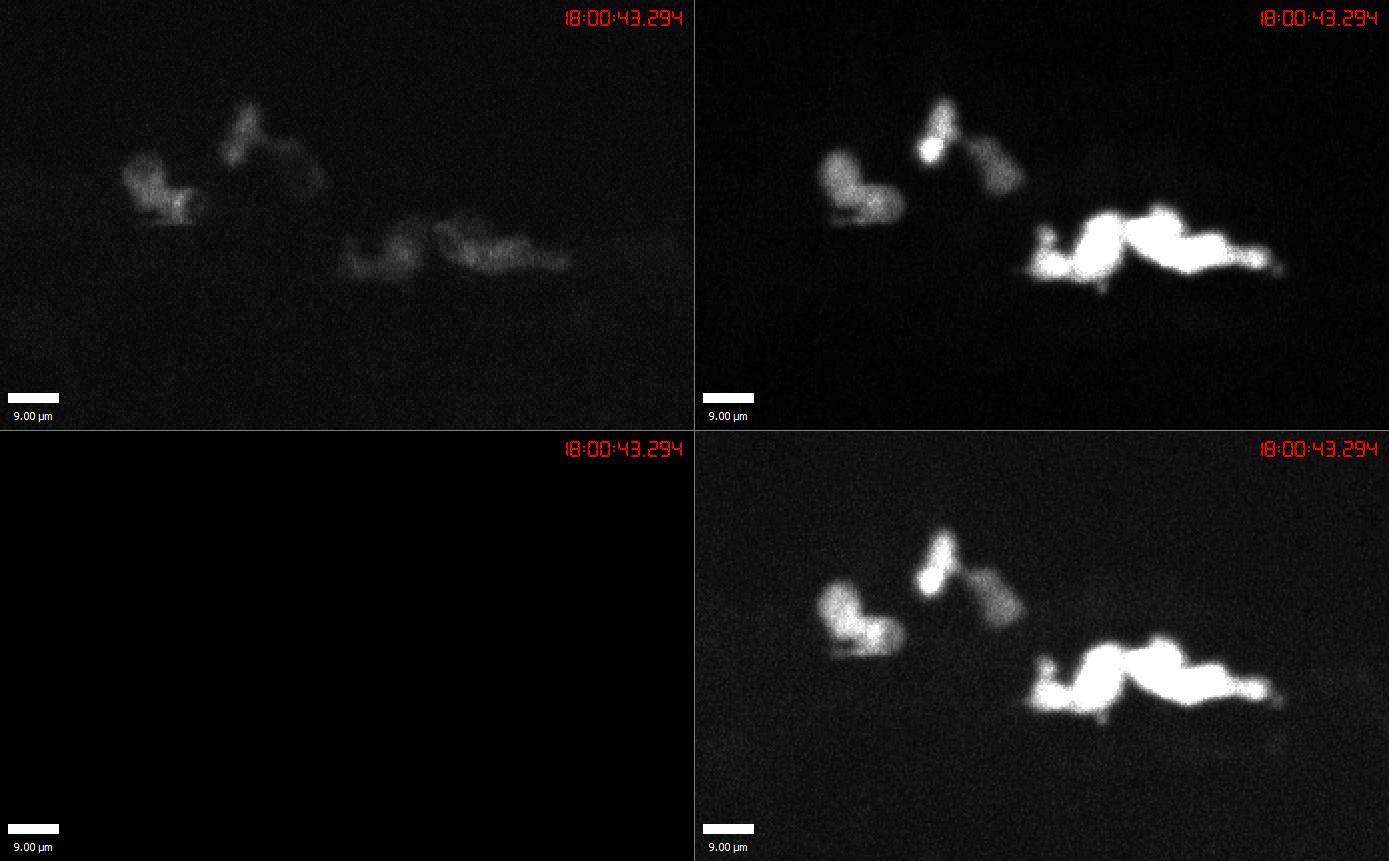

### Het-Aligned-Focused.tif

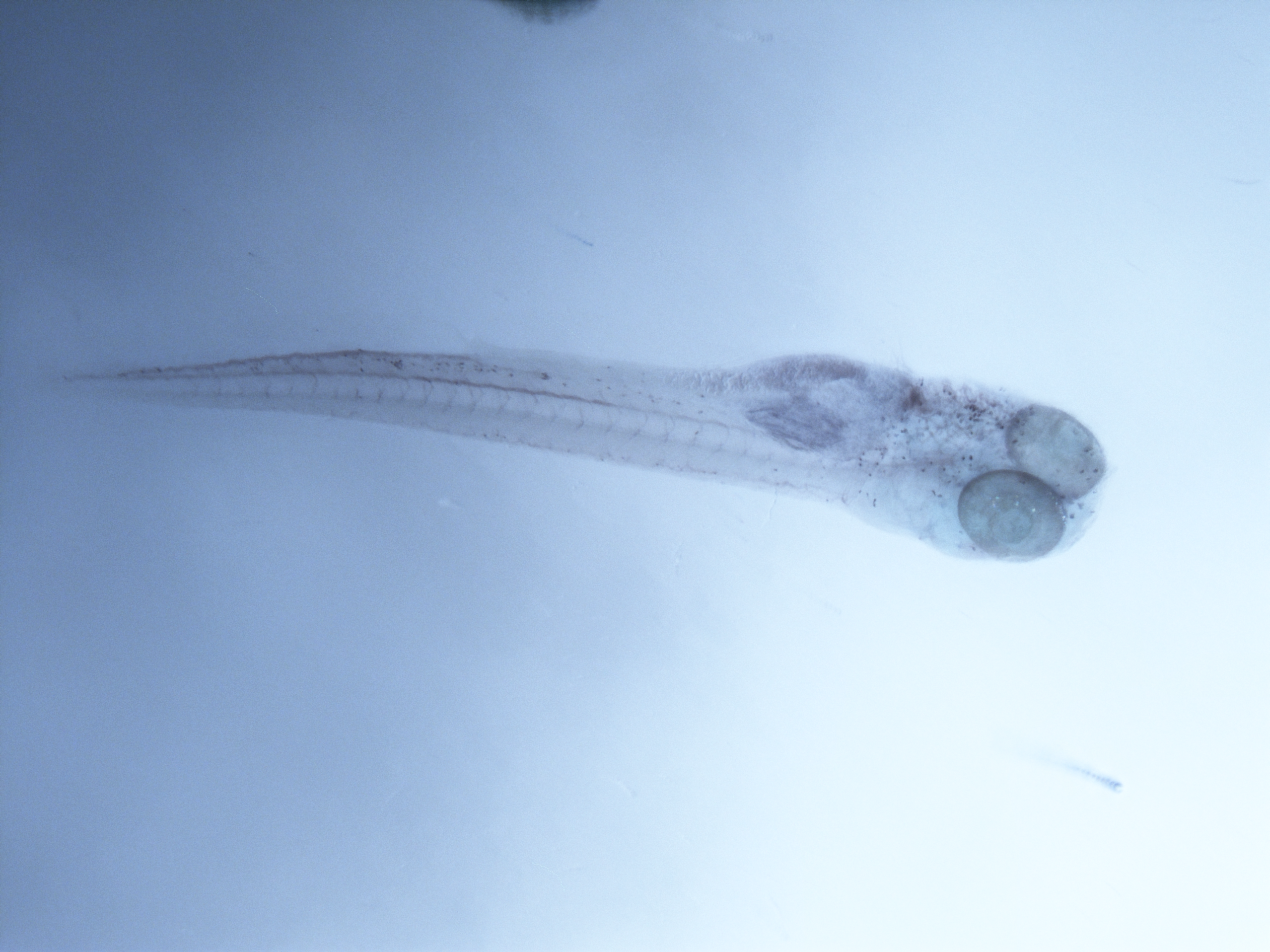

### merged GFP signal.jpg

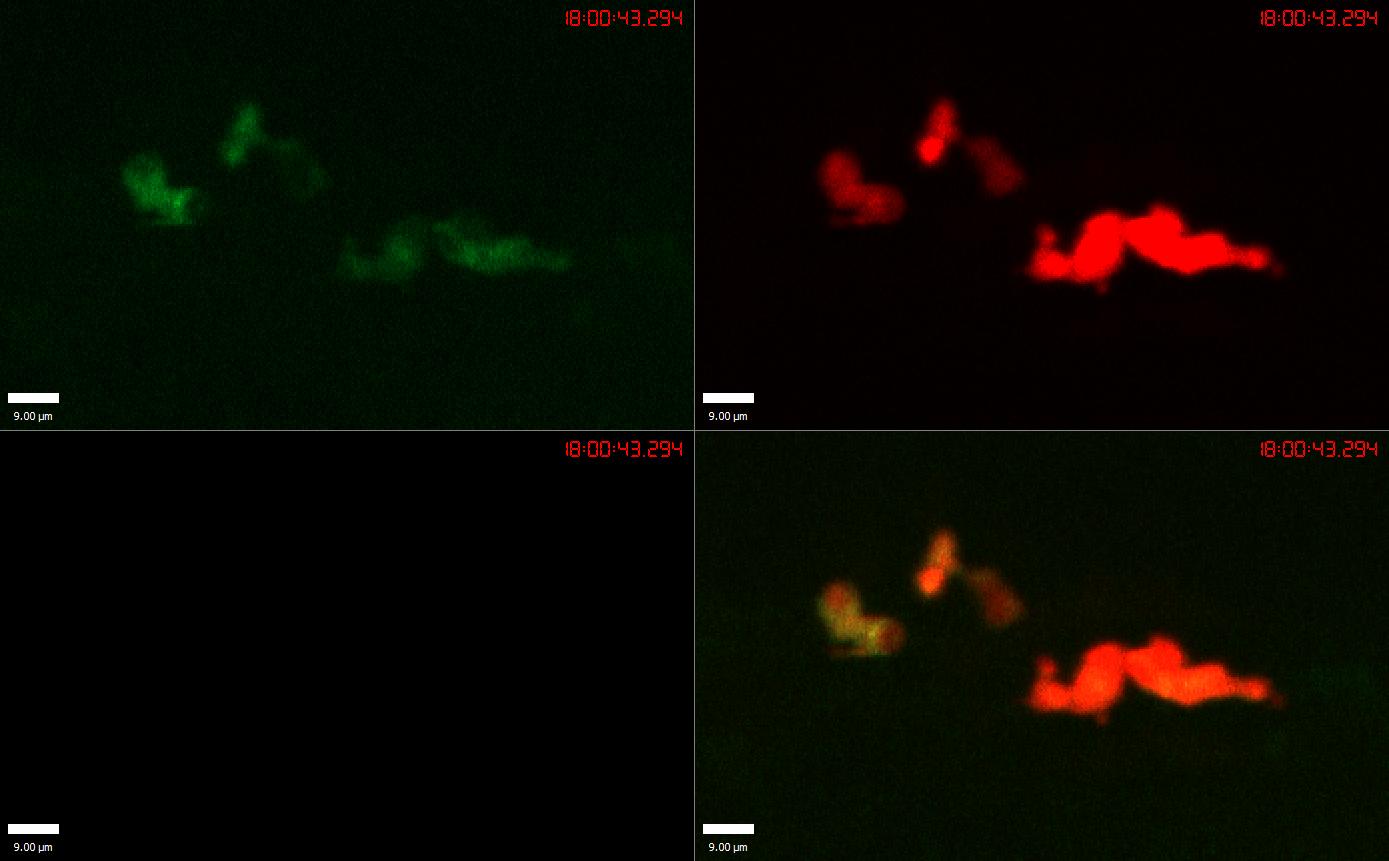

### Sh496 mpx +- x mpx ++ +ve close_crop-Aligned-Focused.tif

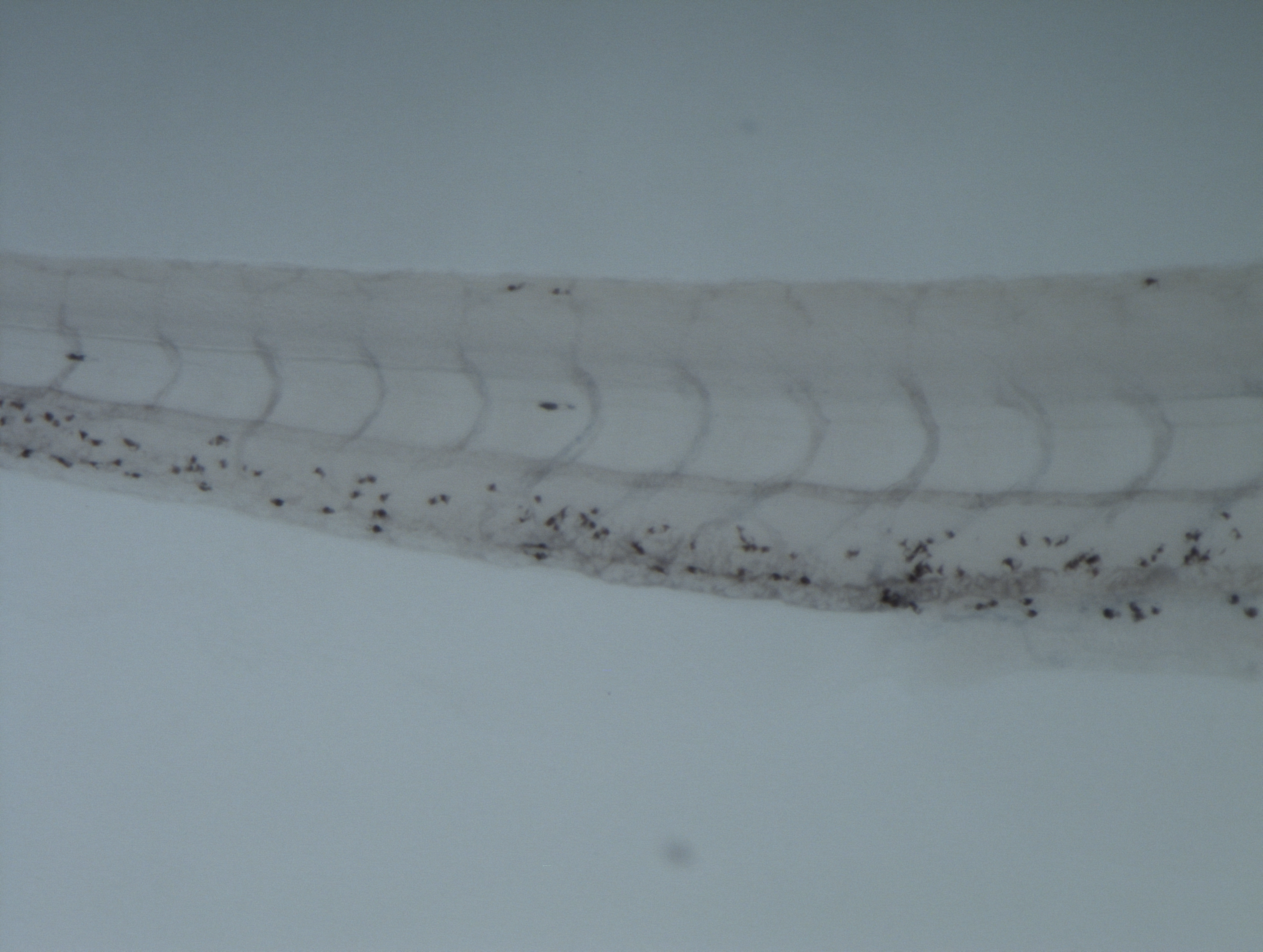

### Sh496 mpx +- x mpx ++ +ve wide_crop-Aligned-Focused.tif

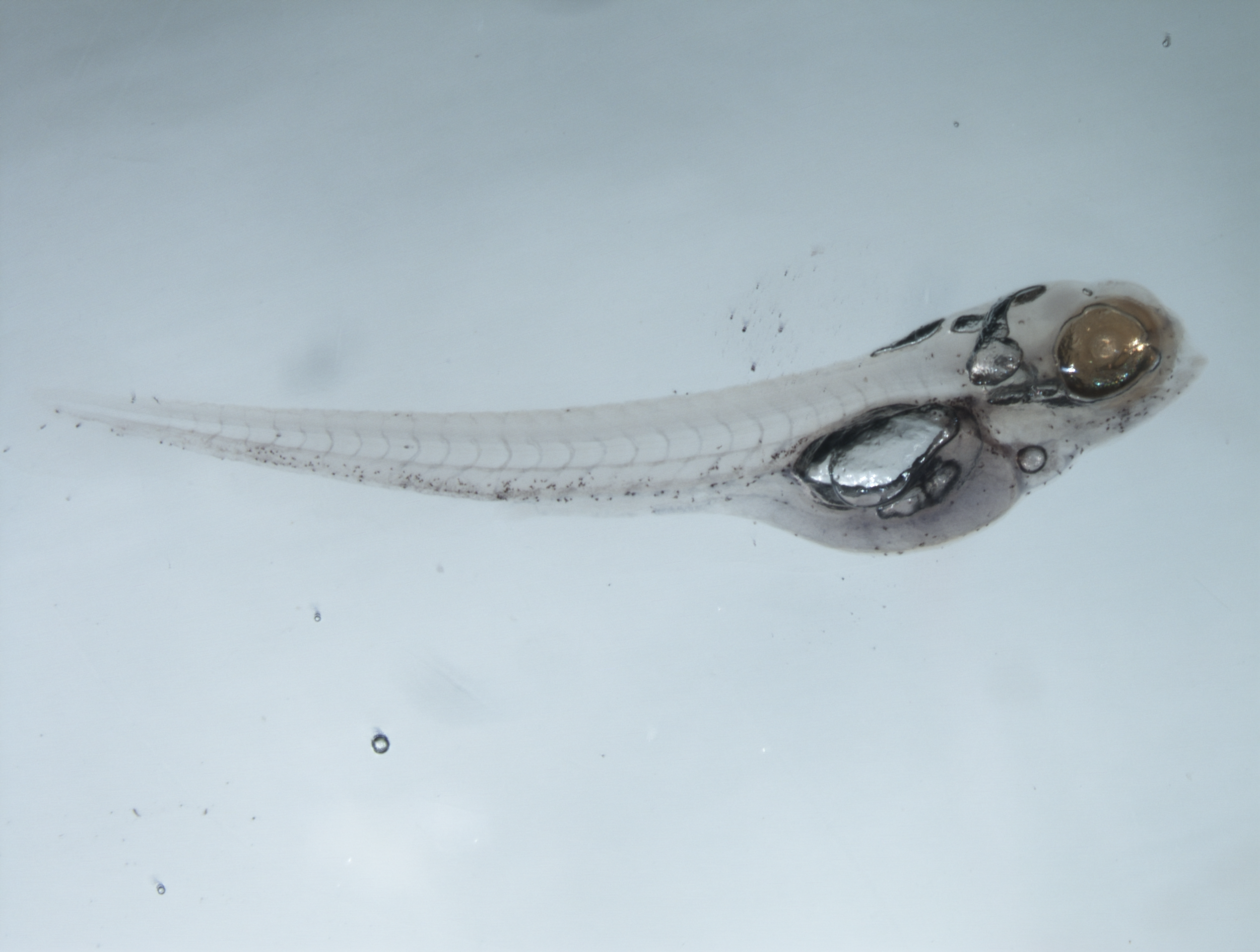

### Sh496 mpx +- x mpx ++ -ve close_crop-Aligned-Focused.tif

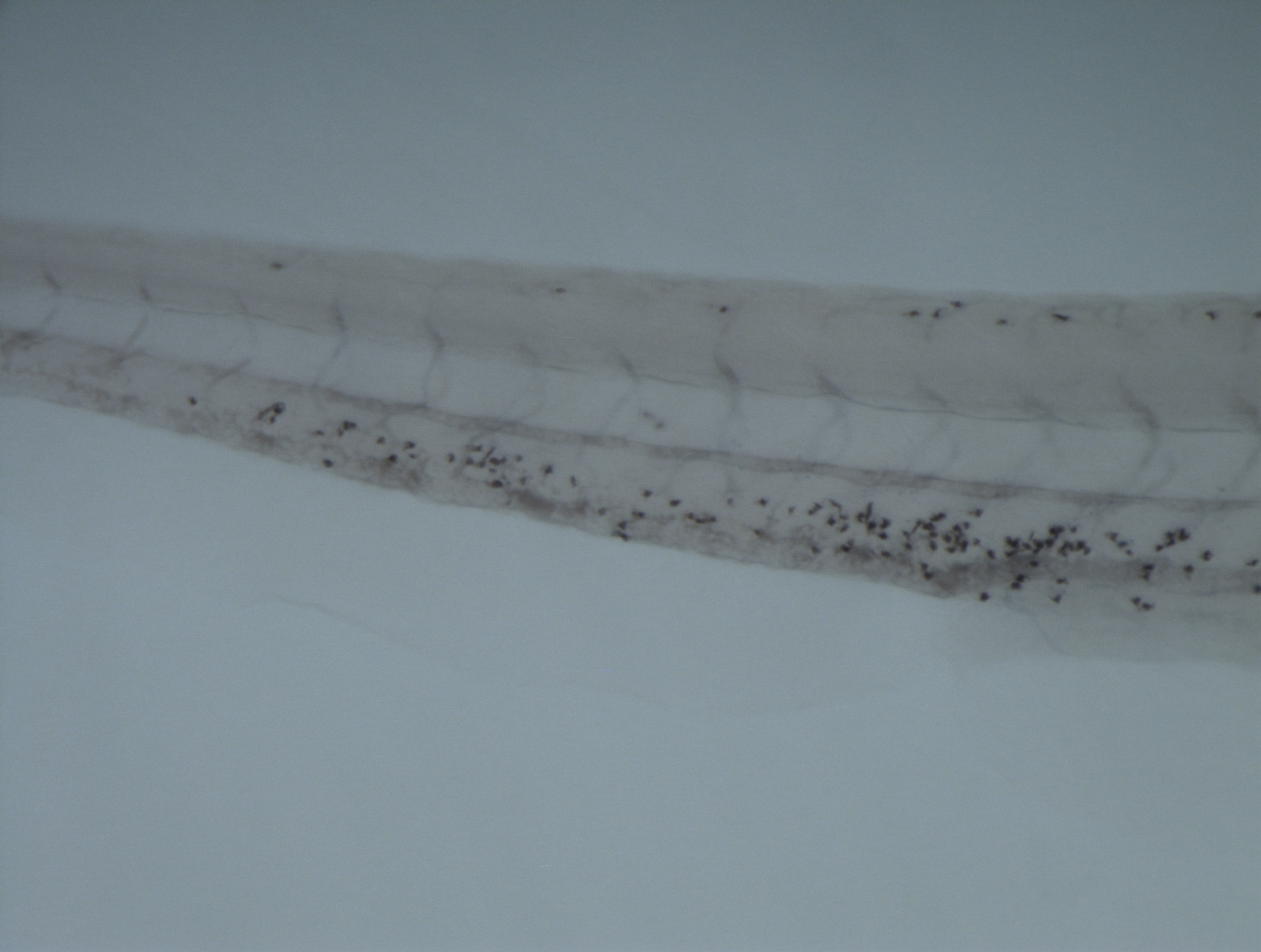

### Sh496 mpx +- x mpx ++ -ve wide_crop-Aligned-Focused.tif

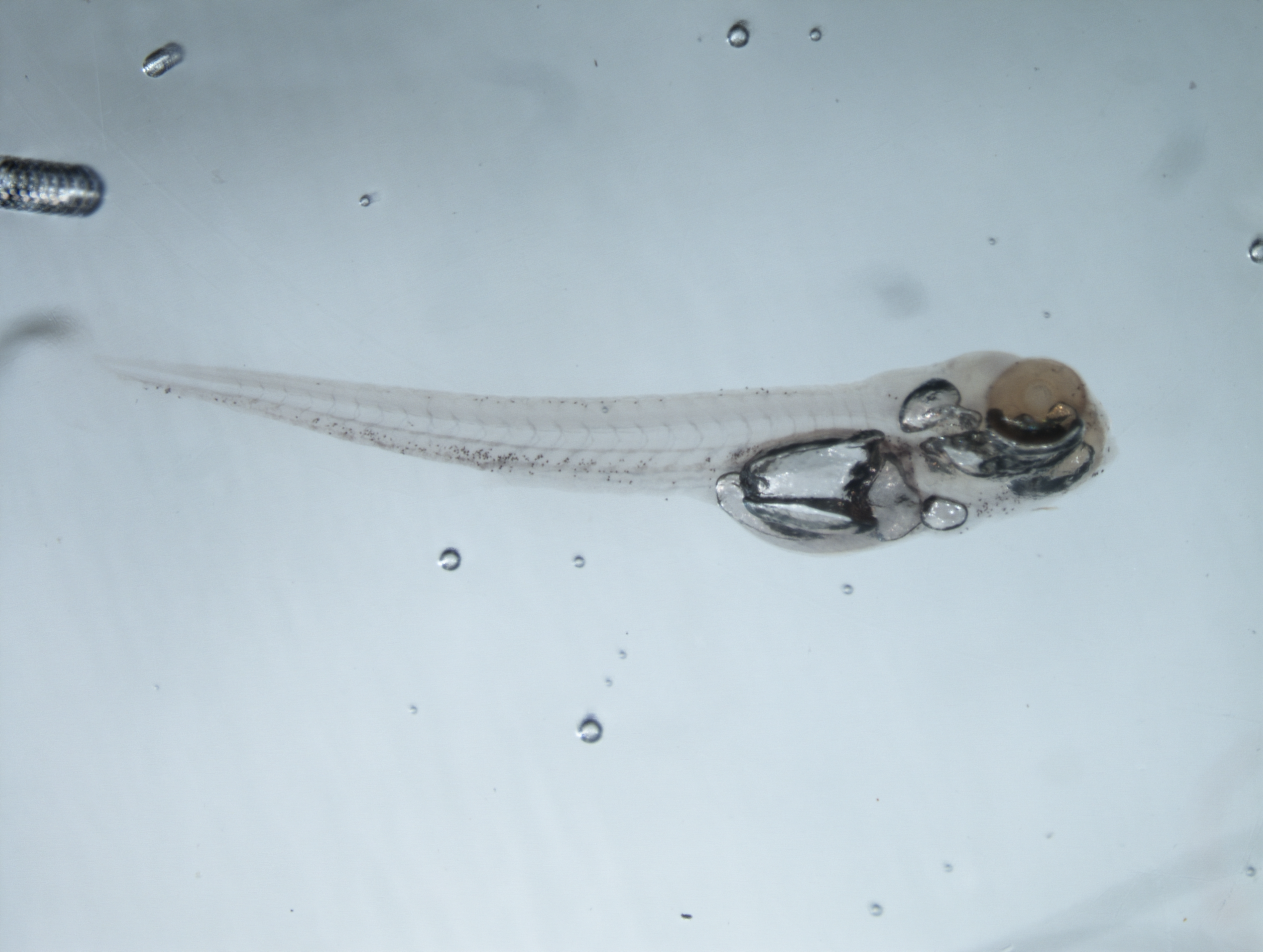

### Sh496 mpx +- x mpx ++ No GH close_crop-Aligned-Focused.tif

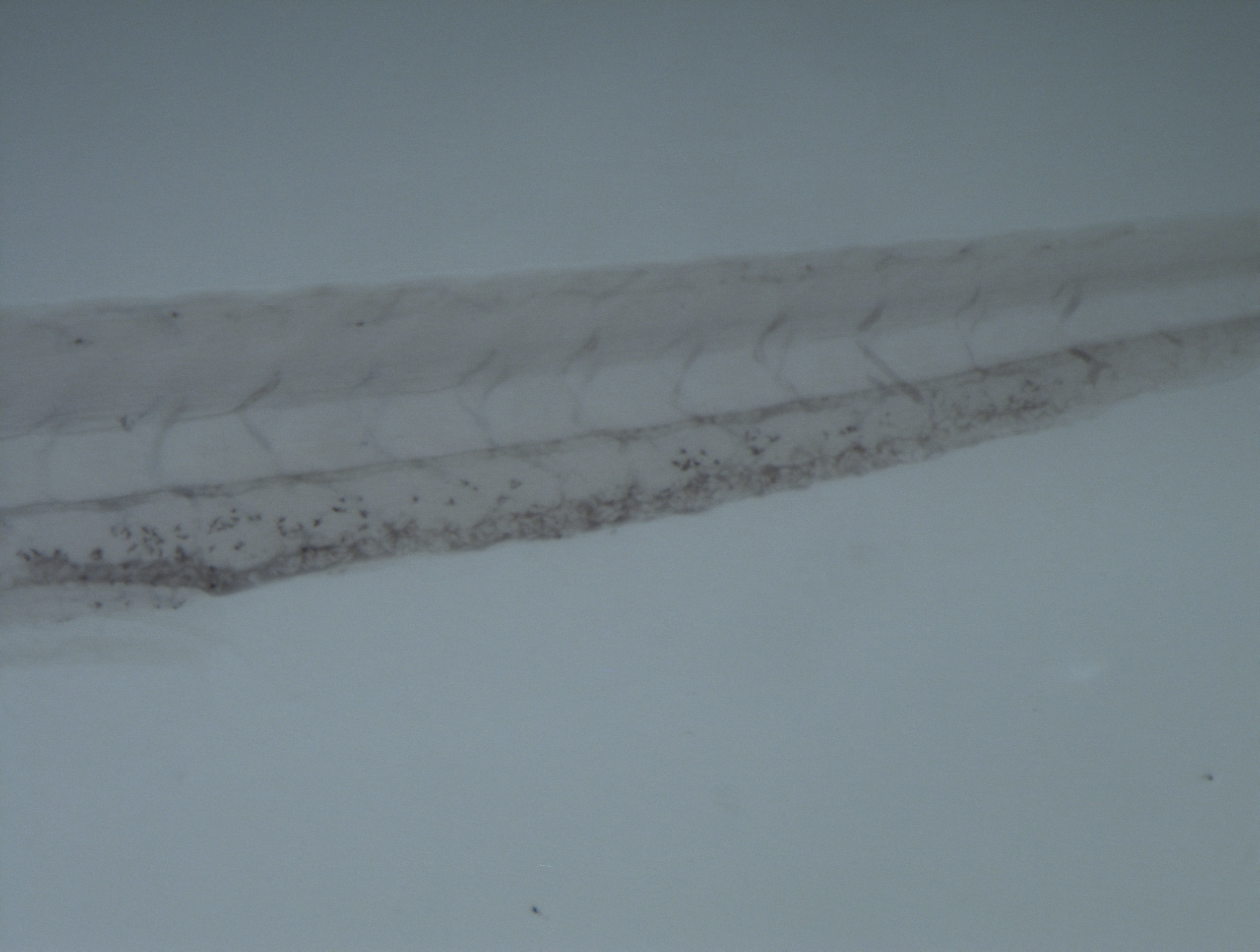

### Sh496 mpx +- x mpx ++ No GH wide_crop-Aligned-Focused.tif

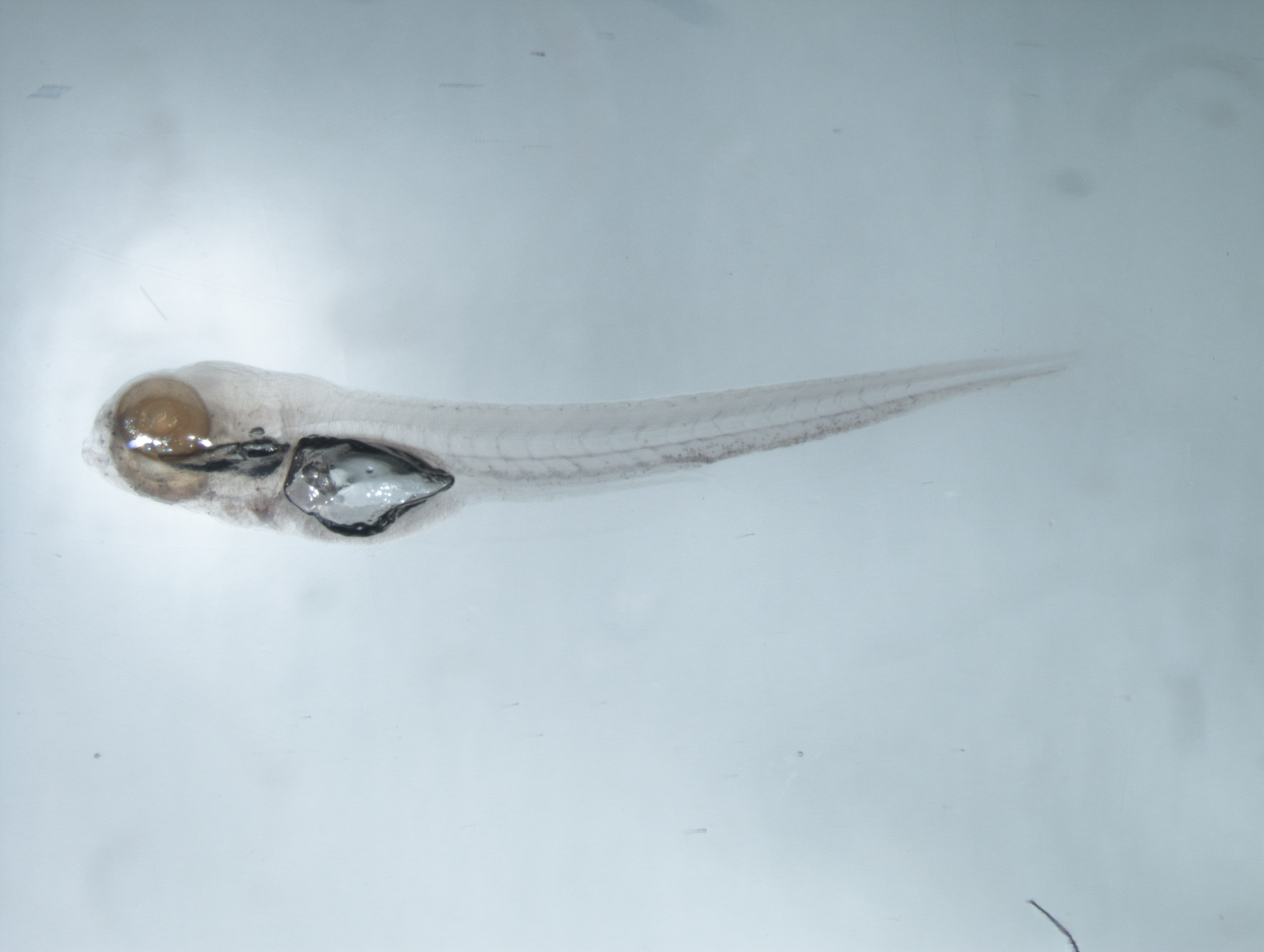

### Sh496 mpx null x mpx null ++ve_crop-Aligned-Focused.tif

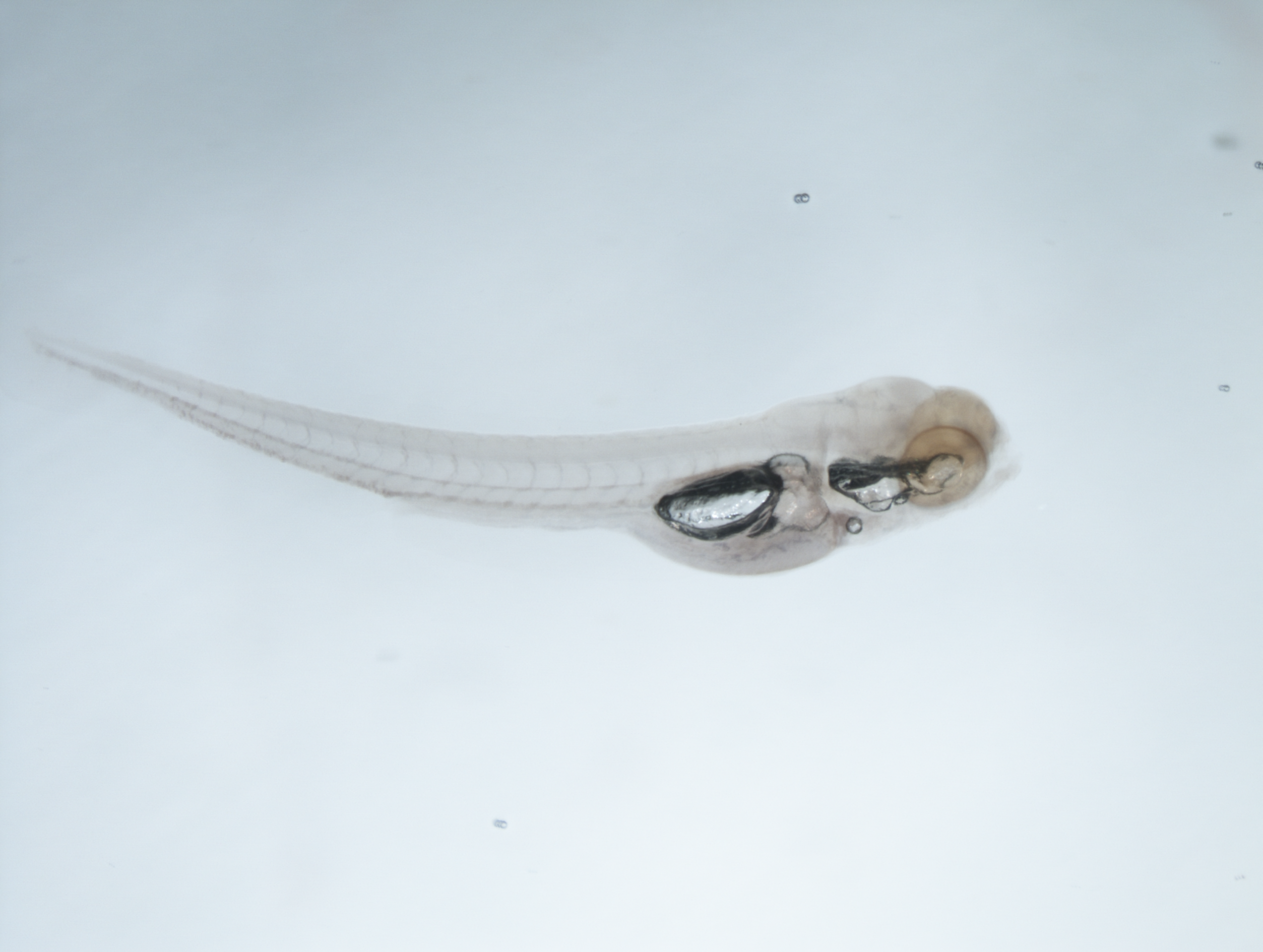

### Sh496 mpx null x mpx null ++veClose_crop-Aligned-Focused.tif

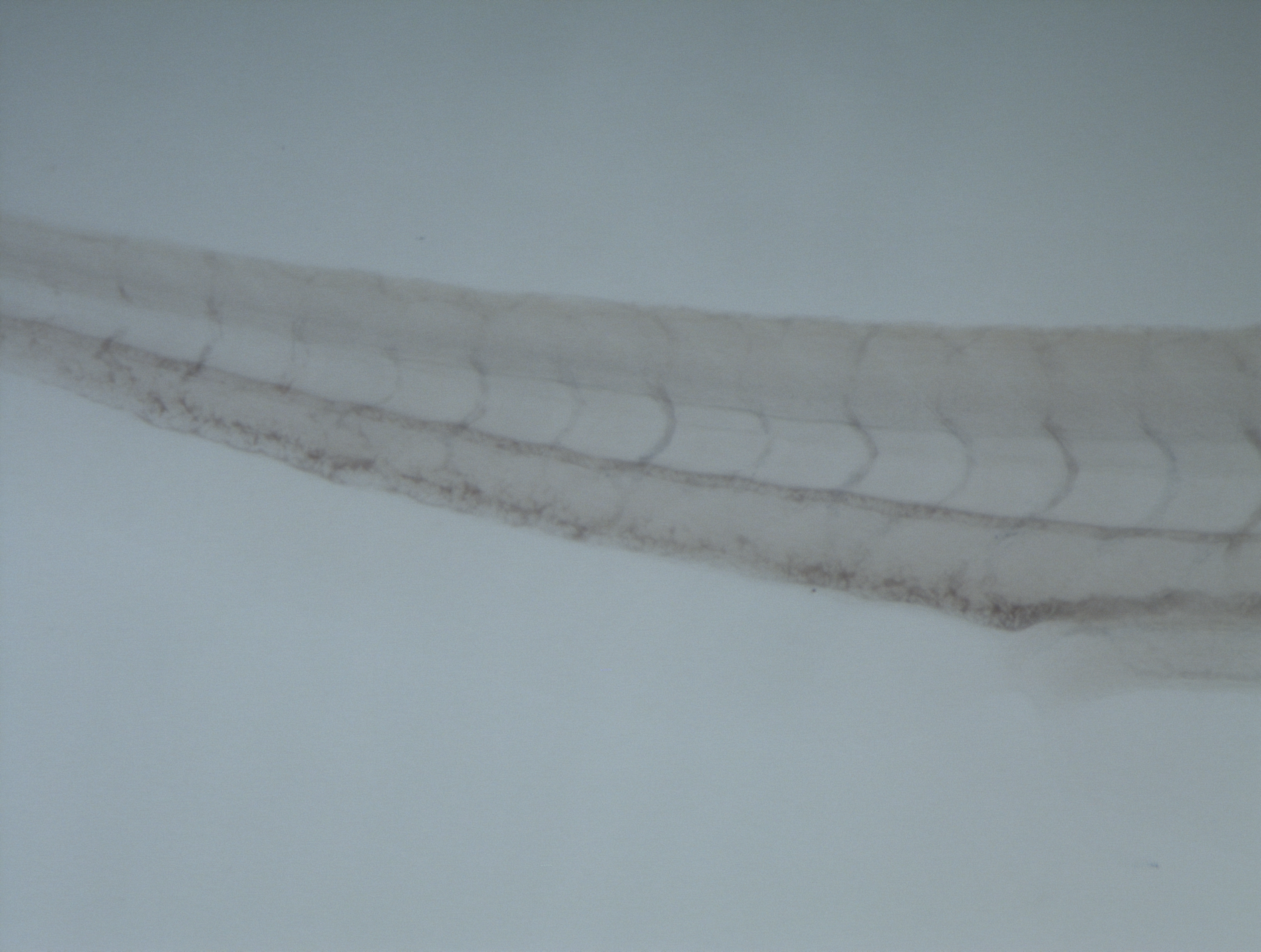

### Sh496 mpx null x mpx null +ve 2 close_crop-Aligned_crop.tif

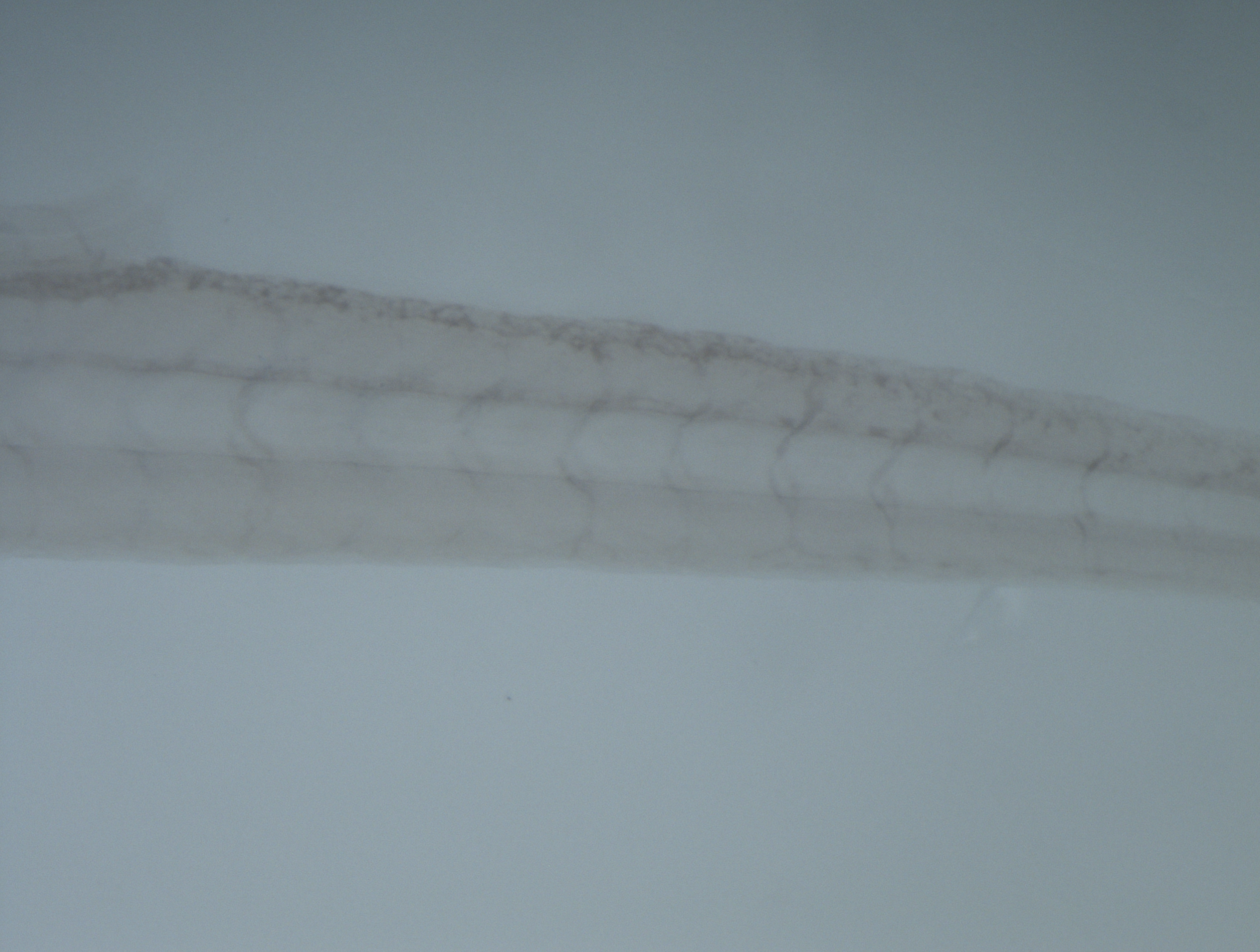

### Sh496 mpx null x mpx null +ve 2 wide_crop.tif

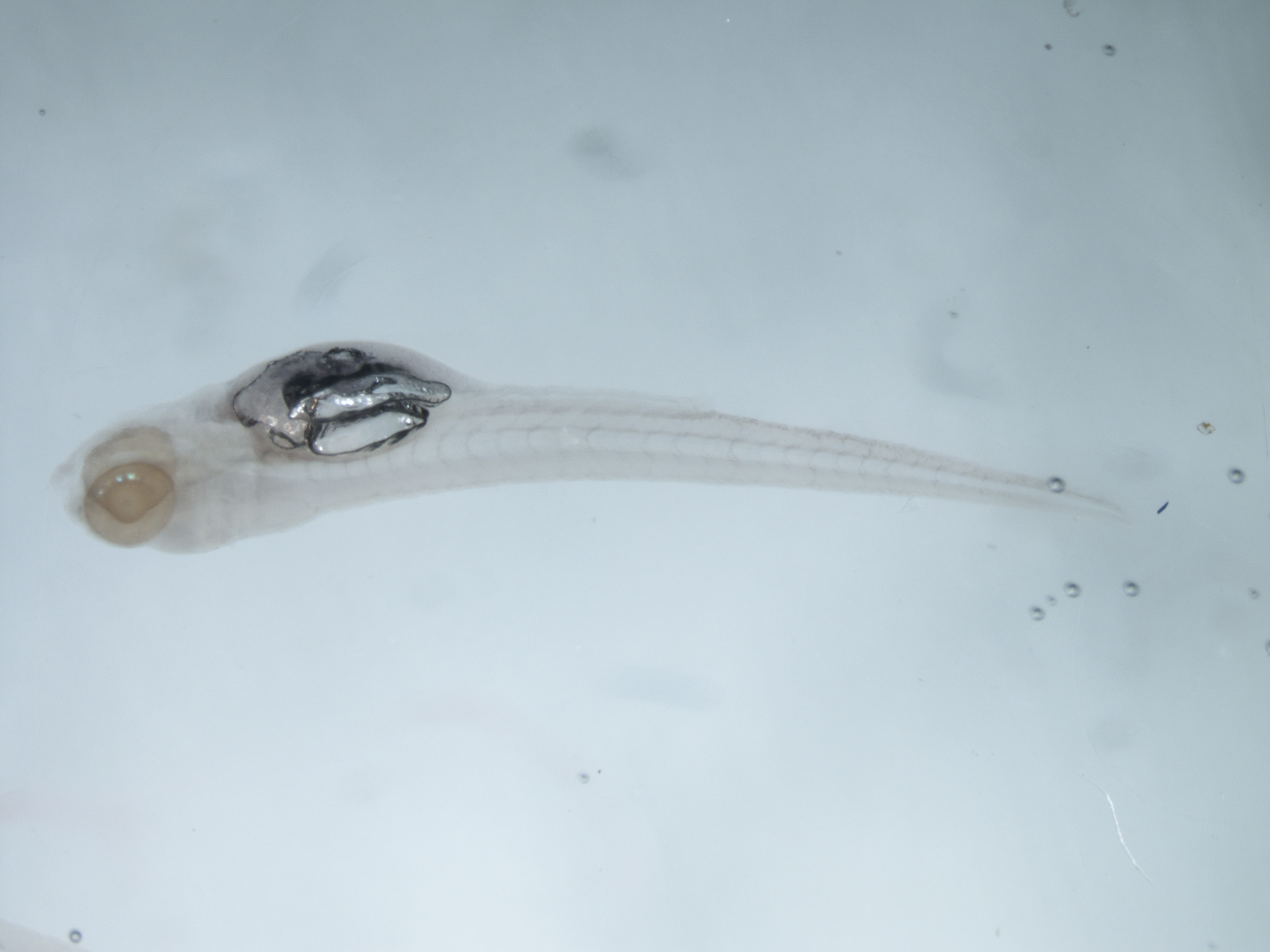

### Sh496 mpx null x mpx null +ve_crop-Aligned-Focused.tif

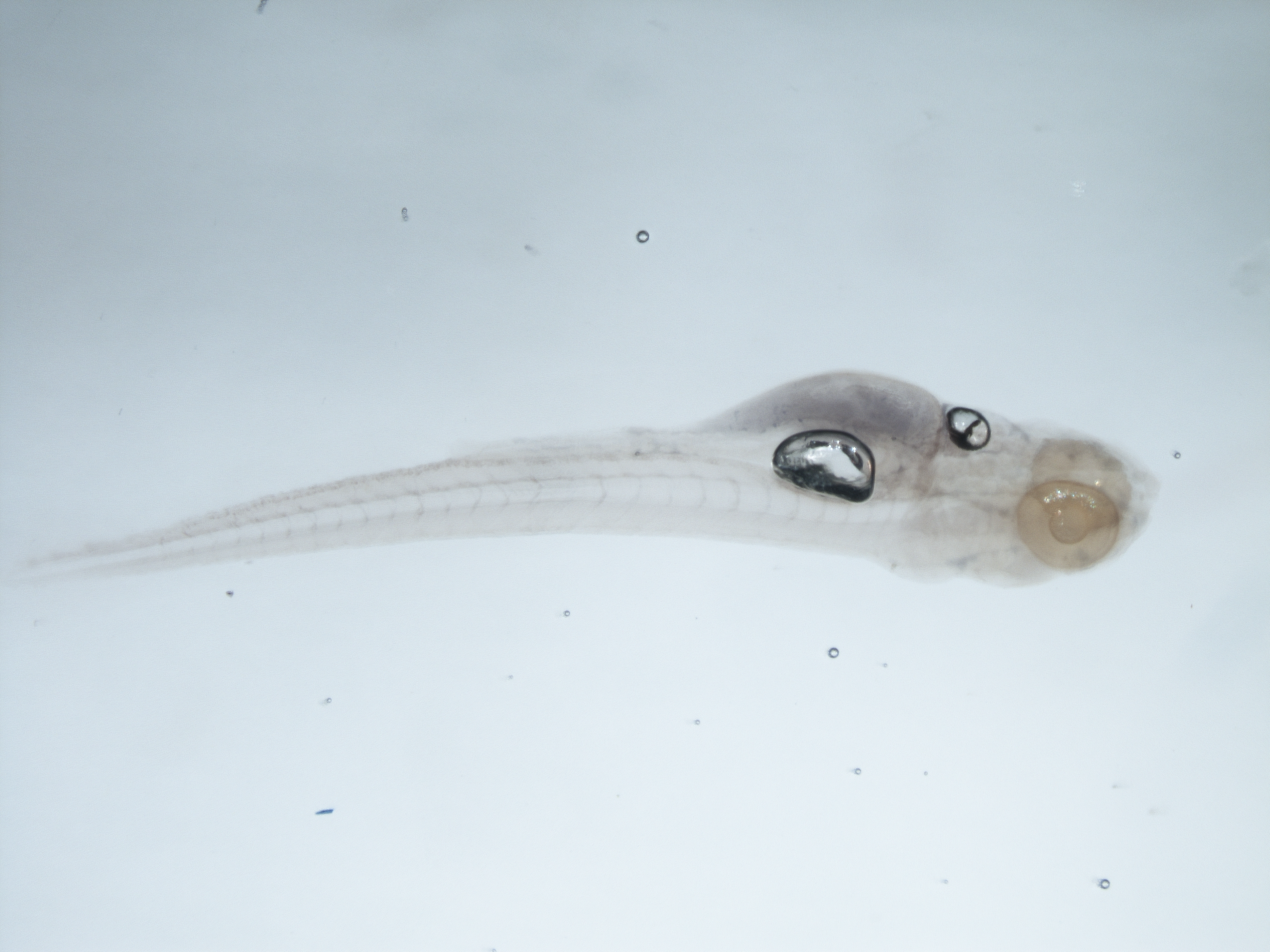

### Sh496 mpx null x mpx null -ve 2 close_crop-Aligned-Focused.tif

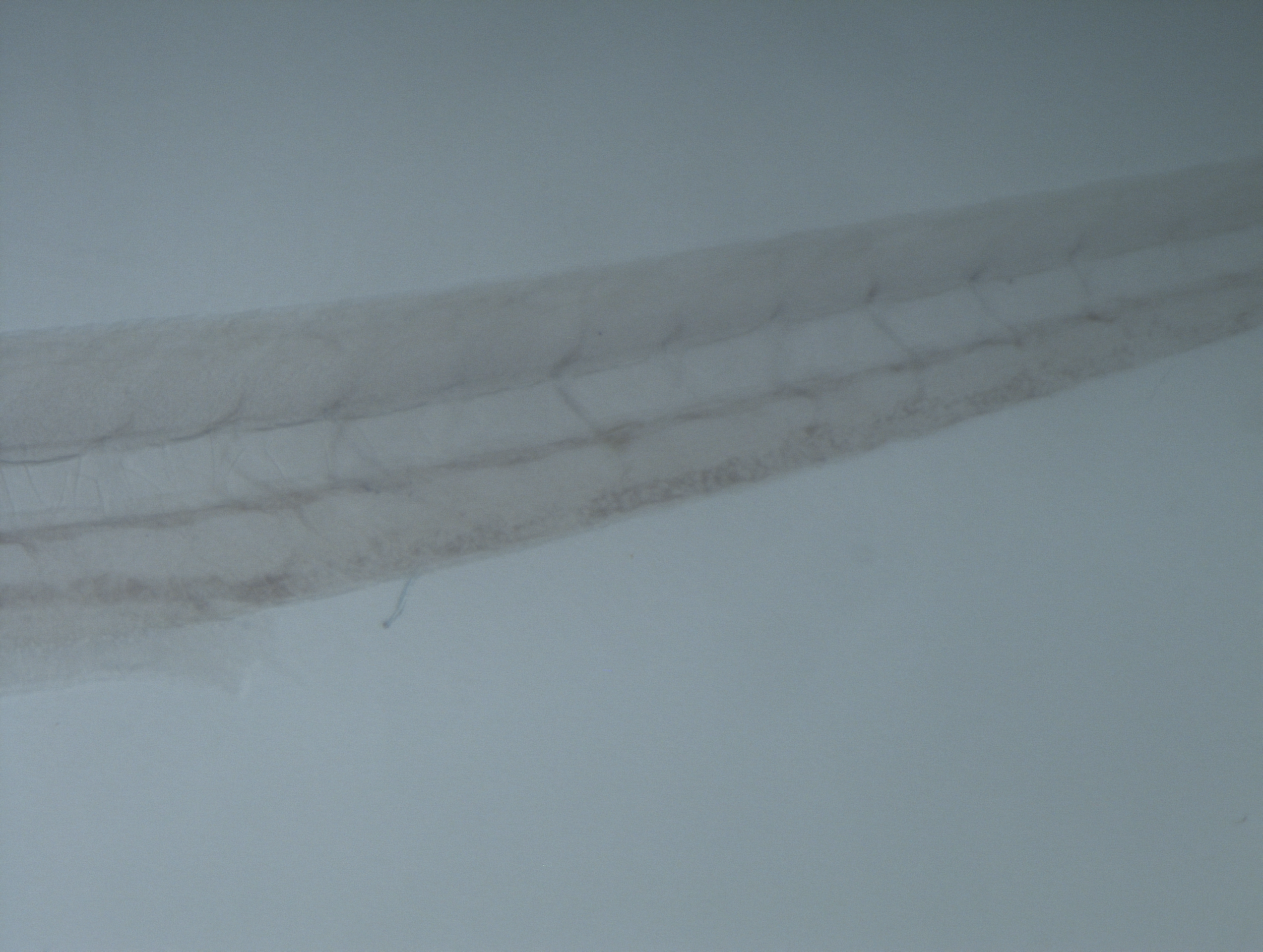

### Sh496 mpx null x mpx null -ve 2 wide_crop.tif

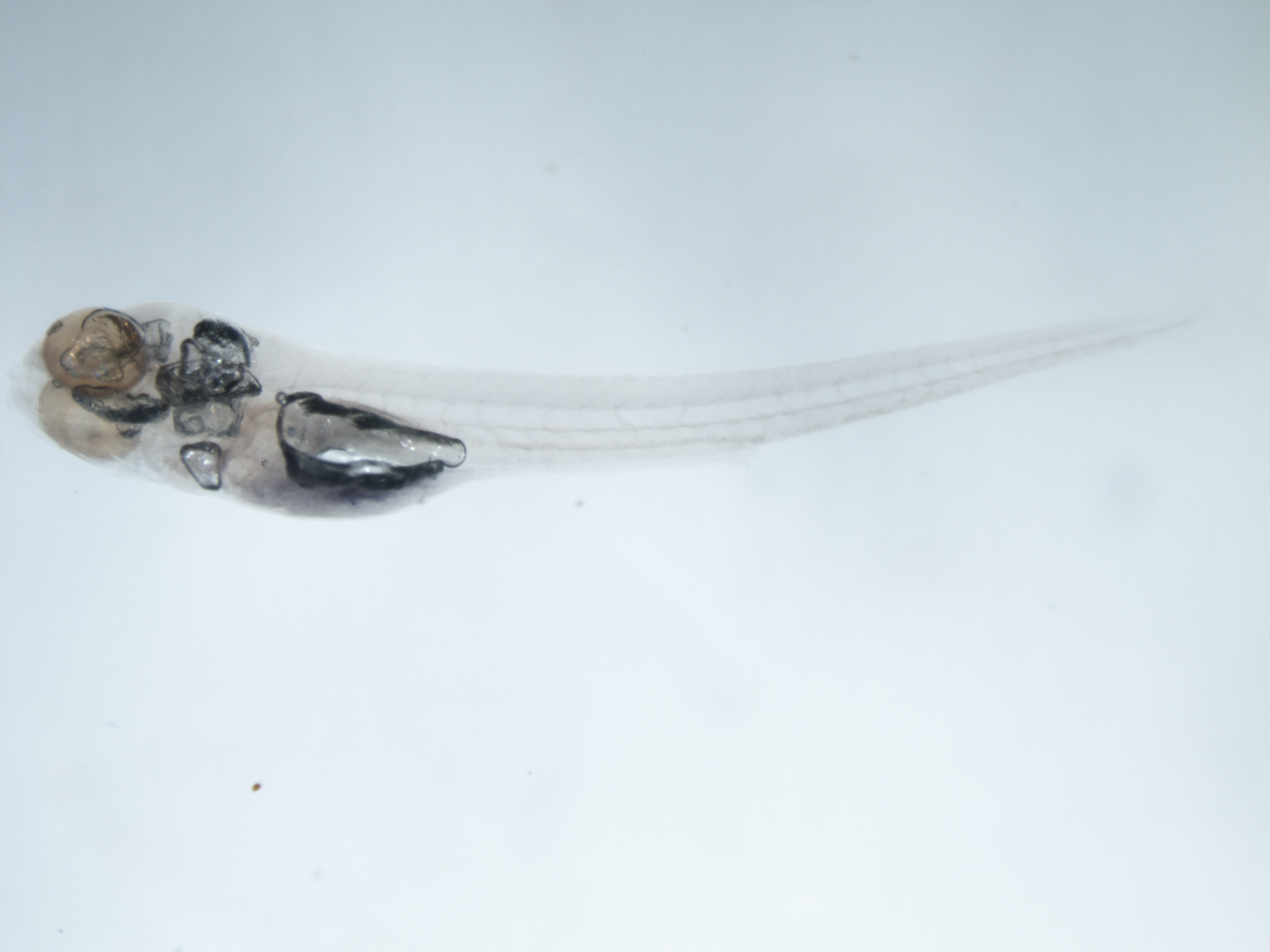

### Sh496 mpx null x mpx null -ve-Aligned_crop.tif

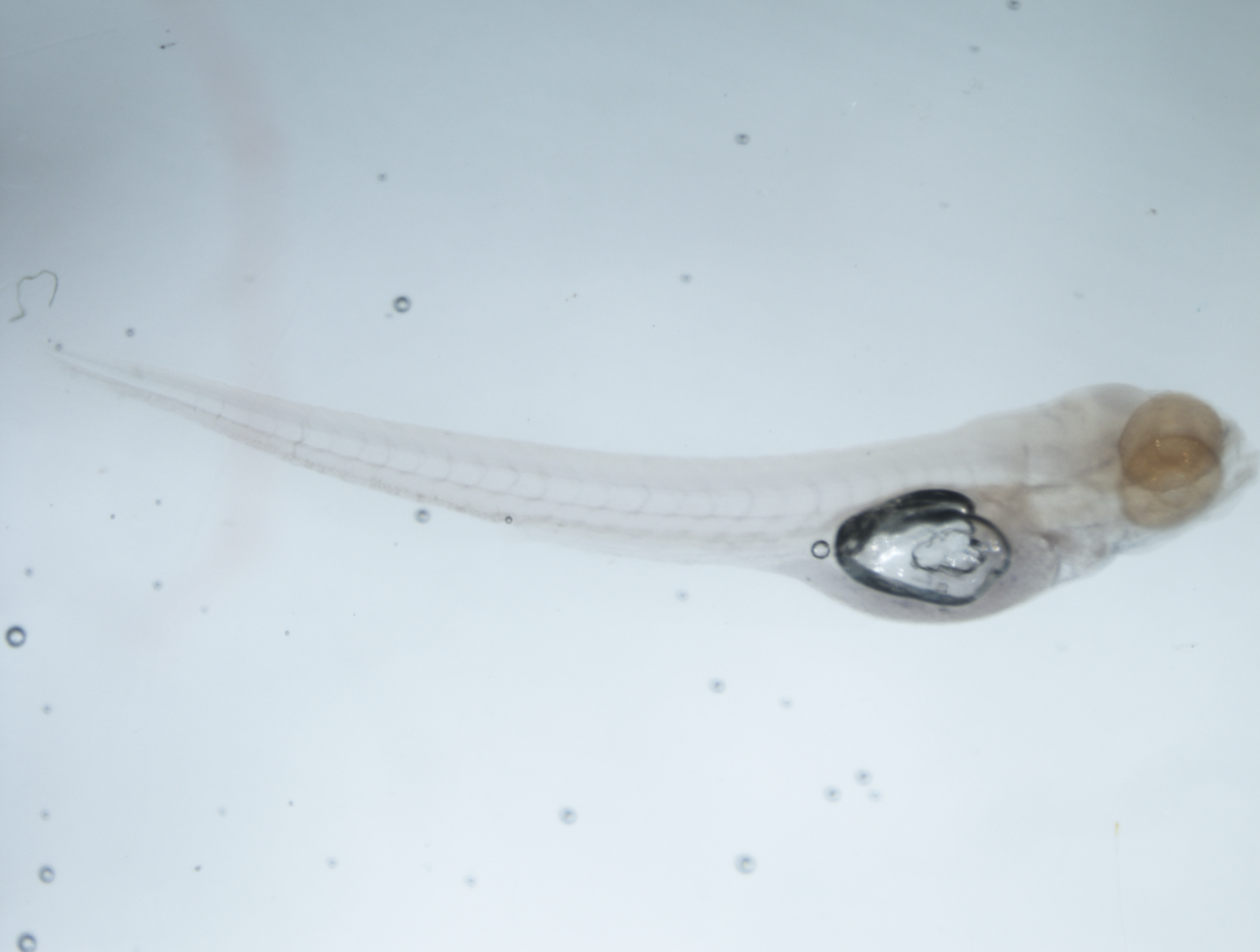

### SH496 Positive close-Aligned-Focused.tif

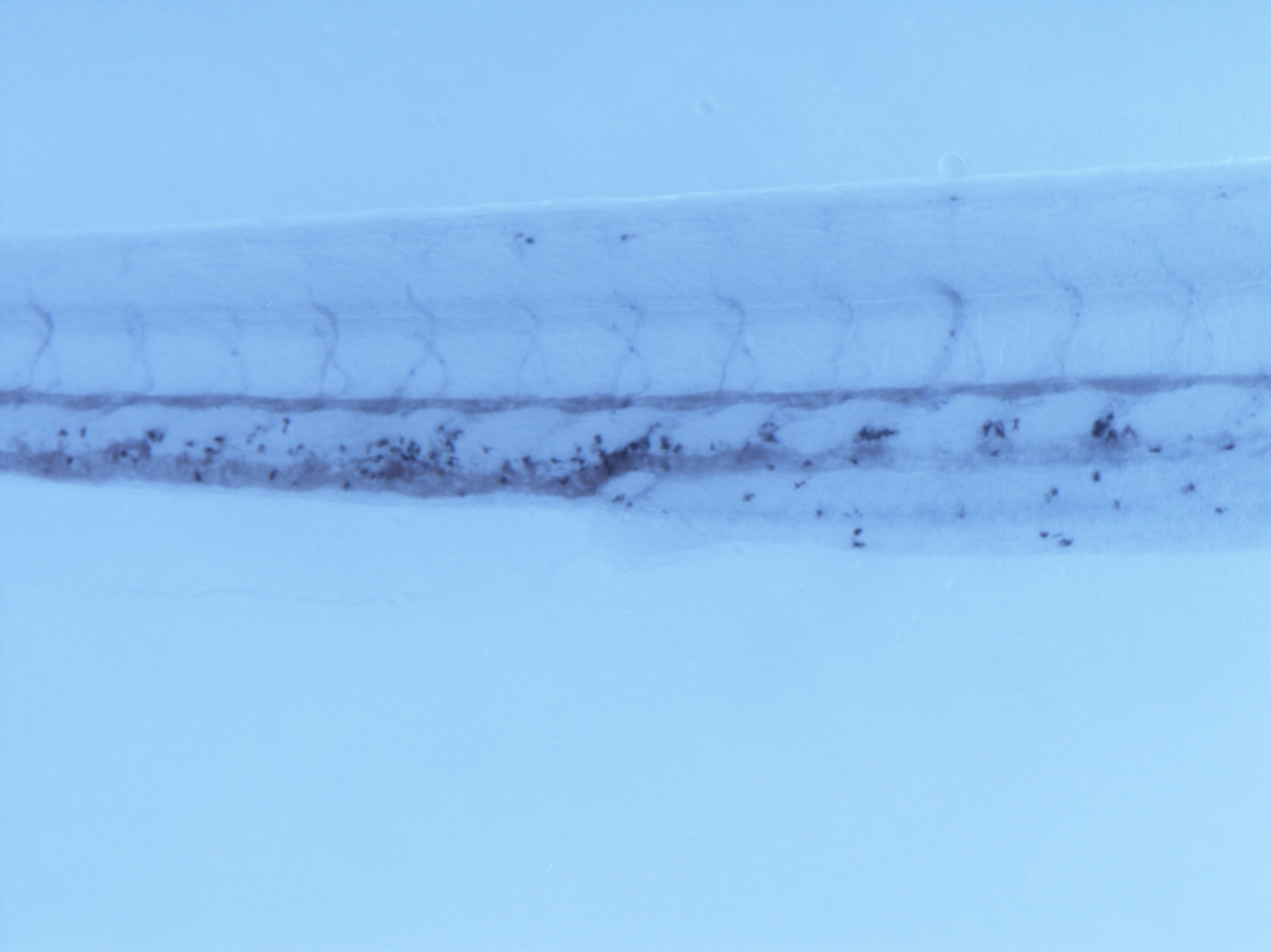

### SH496 Positive-Aligned-Focused.tif

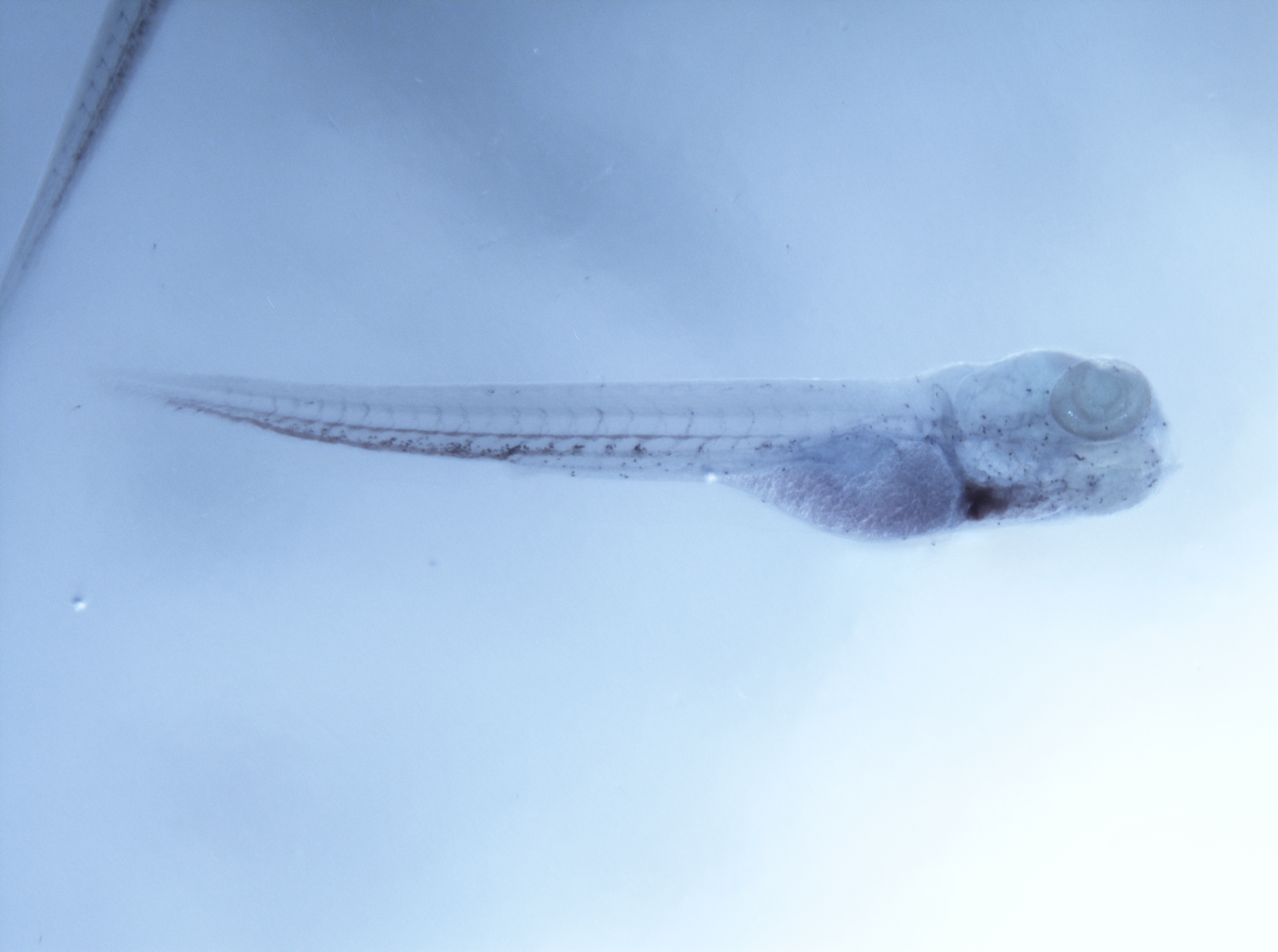

### sh496xsh260 neutrophil Maximum intensity projection.tif

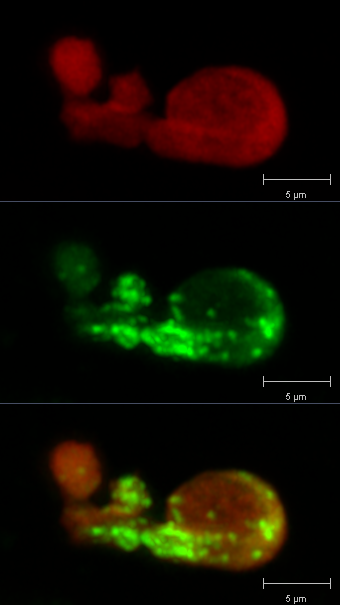

### sh496xsh260 neutrophil zoom final.tif

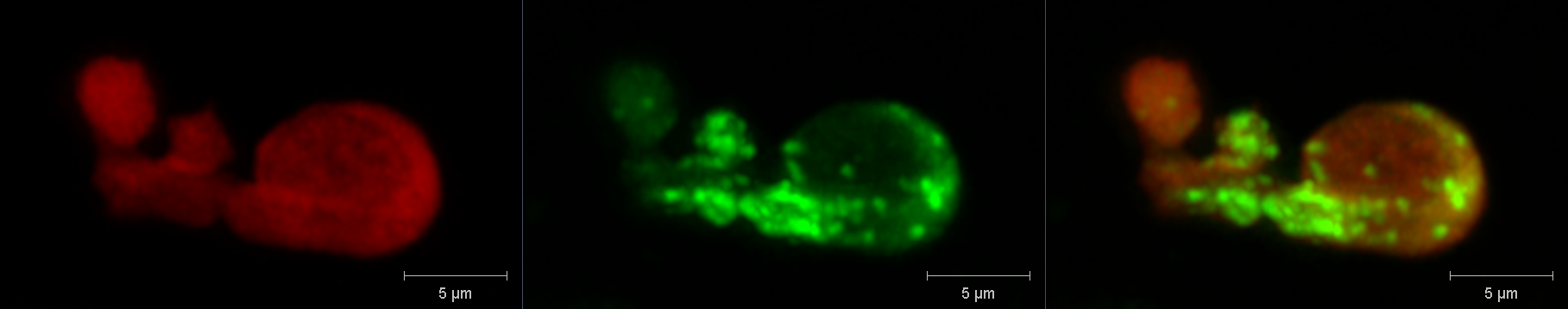

### sh496xsh260 widefield Maximum intensity projection.tif

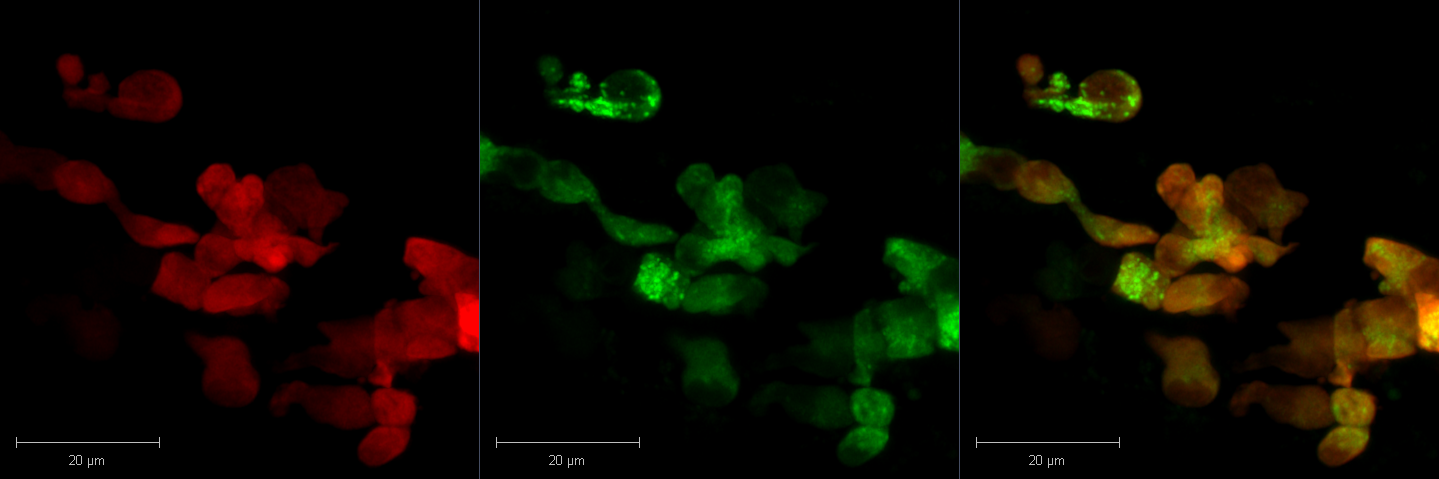

### Spotless close-Aligned_crop.tif

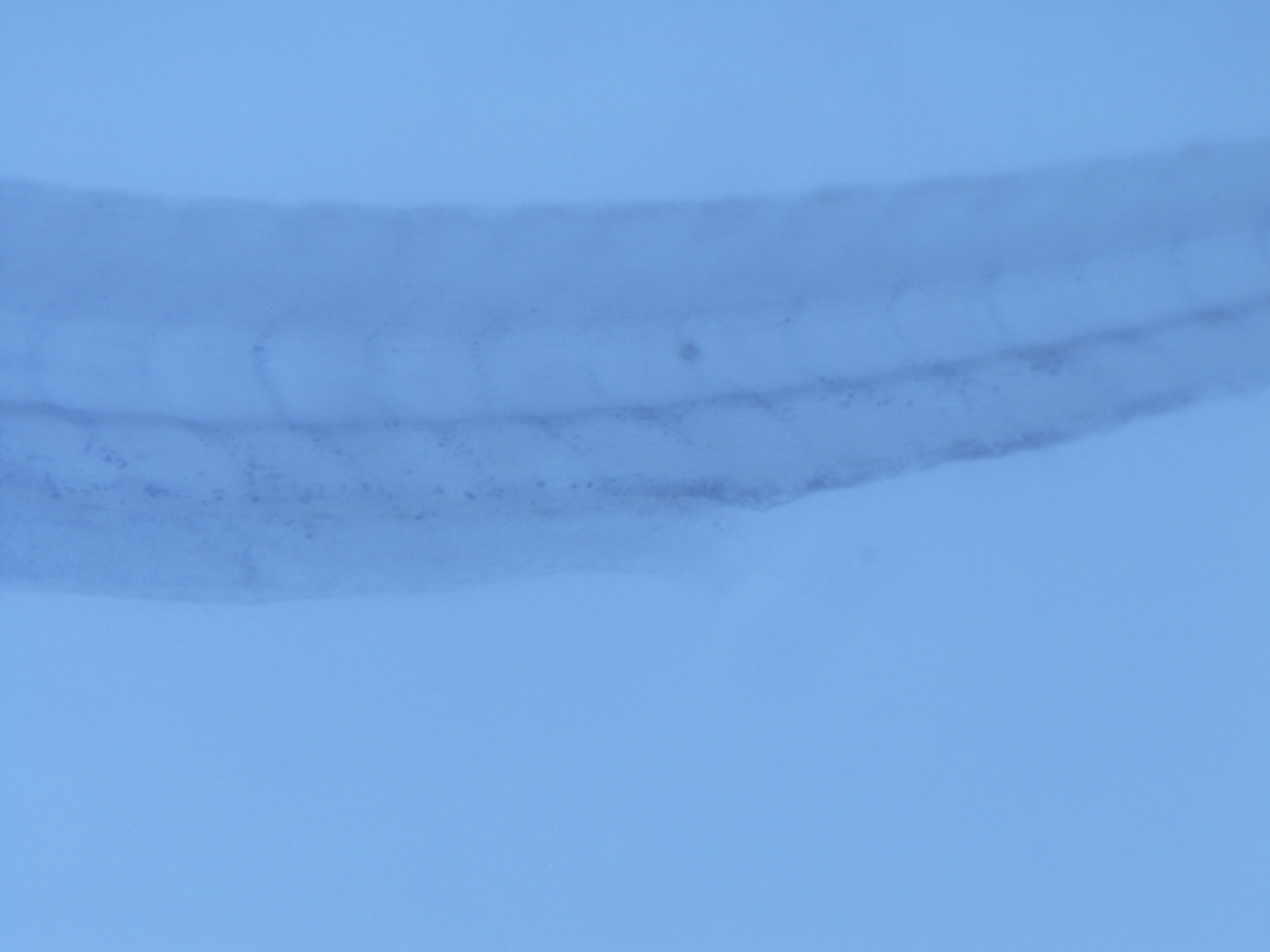

### Spotless Genotyping.tif

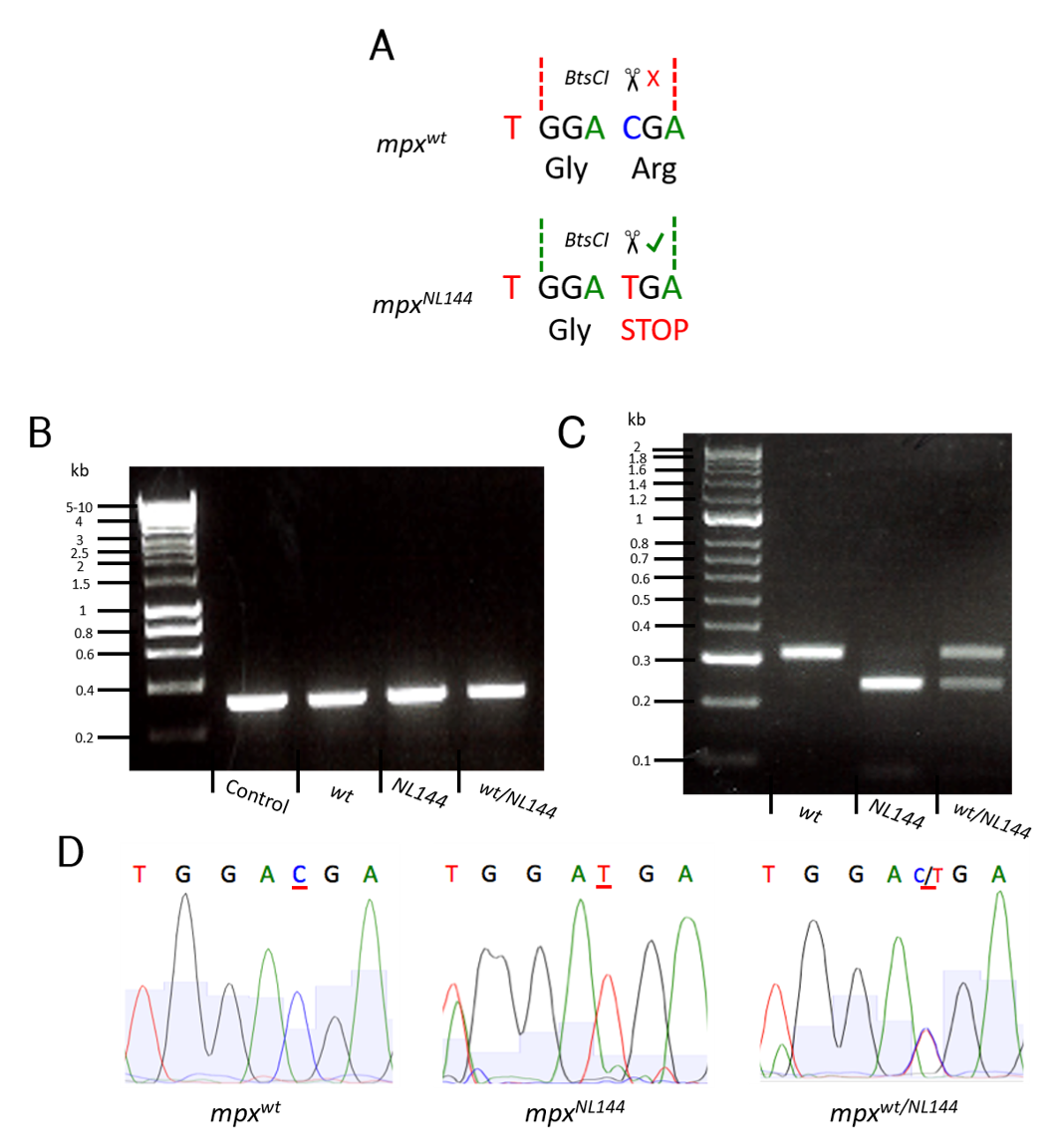

### Spotless-Focused.tif

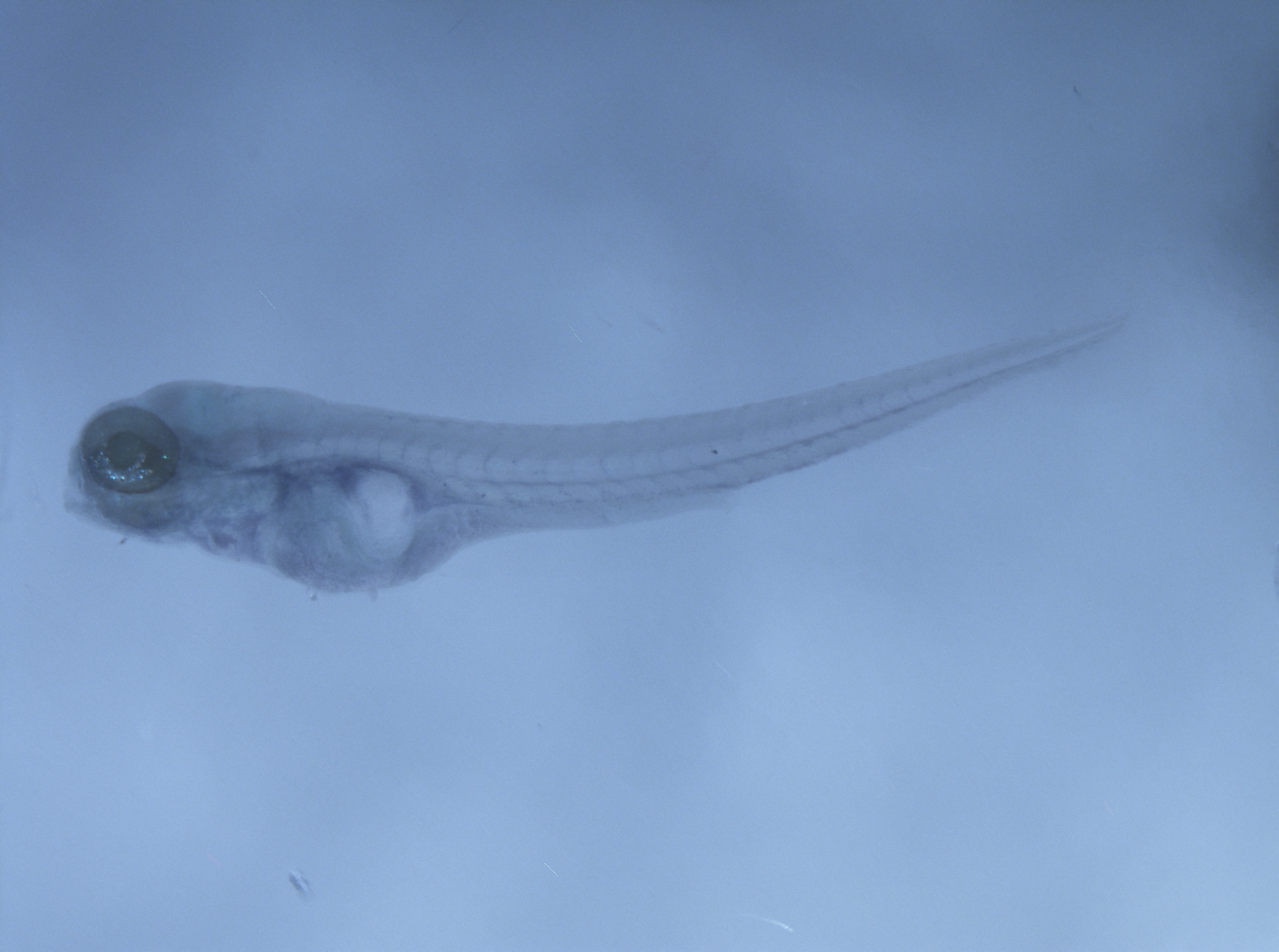
